# Transfer of modified fecal viromes improve blood glucose regulation and alleviates symptoms of metabolic dysfunction-associated fatty liver disease in an obesity male mouse model

**DOI:** 10.1101/2023.03.20.532903

**Authors:** Xiaotian Mao, Sabina Birgitte Larsen, Line Sidsel Fisker Zachariassen, Anders Brunse, Signe Adamberg, Josue Leonardo Castro Mejia, Frej Larsen, Kaarel Adamberg, Dennis Sandris Nielsen, Axel Kornerup Hansen, Camilla Hartmann Friis Hansen, Torben Sølbeck Rasmussen

**Affiliations:** Section of Food Microbiology, Gut Health, and Fermentation, Dept. of Food Science, University of Copenhagen, Frederiksberg, Denmark; Section of Experimental Animal Models, Dept. of Veterinary and Animal Sciences, University of Copenhagen, Frederiksberg, Denmark; Section of Comparative Pediatrics and Nutrition, Dept. of Veterinary and Animal Sciences, University of Copenhagen, Frederiksberg, Denmark; Department of Chemistry and Biotechnology, Tallinn University of Technology, Tallinn, Estonia

**Author notes:** Address correspondence to, Rolighedsvej 26 4th floor, 1958 Frederiksberg C, Denmark.

**Keywords:** Fecal virome transplantation, gut microbiome, type-2 diabetes, MAFLD, phage therapy, obesity

## Abstract

Metabolic syndrome encompasses amongst other conditions like obesity, type-2 diabetes, and metabolic dysfunction associated fatty liver disease (MAFLD), which are all associated with gut microbiome (GM) dysbiosis. Fecal microbiota transplantation (FMT) has been explored to treat metabolic syndrome by restoring the GM. FMT is generally safe, but motivated by case reports, accidental transfer of pathogenic bacteria remains a concern. As a safer alternative, fecal virome transplantation (FVT, sterile-filtrated feces) has the advantage over FMT in that mainly bacteriophages are transferred and FVT from lean male donors has shown promise in alleviating the metabolic effects of a high-fat diet in a preclinical mouse study. However, FVT still carries the risk of eukaryotic viral infections. To address this, we here apply recently developed modification methodologies to inactivate or remove the eukaryotic viral component of FVT while maintaining an active enteric bacteriophage community. Modified FVTs were compared with unmodified FVT and saline in an animal model of diet-induced obesity using male C57BL/6N mice. In contrast to the obese control group, mice administered a modified FVT, nearly depleted from eukaryotic viruses (0.1%), exhibited enhanced blood glucose clearance, although without a concurrent reduction in weight gain. The unmodified FVT improved liver pathology and reduced the proportions of immune cells in the adipose tissue with a non-uniform response. GM analysis suggested that bacteriophage-mediated GM modulation had influenced these outcomes. When optimized, this may pave the way for developing safe bacteriophage-based therapies targeting metabolic syndrome through GM restoration.

## Introduction

Obesity is a severe metabolic disease that affects millions of individuals worldwide due to high-calorie foods and a sedentary lifestyle^1^. It is also a risk factor for the later development of other metabolic comorbidities, such as type-2 diabetes and metabolic dysfunction-associated fatty liver disease (MAFLD)^2,3^. While many patients with obesity can achieve significant weight loss, only a few can maintain their long-term body weight. It is increasingly recognized that a dysbiotic gut microbiome (GM) plays a crucial role in the persistence of obesity^2,4^. Therefore, targeting the GM for treating metabolic syndrome and restoring its long-term balance is an attractive strategy. Fecal microbiota transplantation (FMT) has shown high treatment efficacy in targeting recurrent *Clostridioides difficile* infections (rCDI)^5^, as well as potential importance in colorectal cancer treatments^6^, and the prevention of necrotizing enterocolitis^7,8^. FMT has also been investigated to alleviate symptoms of metabolic syndrome. Although some positive effects have been reported when using FMT to treat patients suffering from obesity and type-2 diabetes^9–11^, these effects are usually limited and short-term. Despite the high success rates in treating rCDI^5^ and the rigorous donor screening, FMT remains a last resort for patients who do not respond to antibiotics due to safety concerns^12^. The reasoning for this caution was exemplified by the death of a patient in 2019 due to a bacterial infection following FMT^13^, as well as subsequent safety alerts from the authorities^14,15^. Consequently, while FMT has the potential to revolutionize treatments for many gut-related diseases, its widespread use seems implausible due to the inherent safety issues of transferring pathogenic microorganisms. The fecal matrix of FMT includes bacteria, archaea, eukaryotes, viruses, and gut metabolites, which may be responsible for both the beneficial and detrimental effects of FMT^16^. Therefore, reducing the complexity of the donor fecal matrix while maintaining its therapeutic efficacy would improve safety in clinical settings.

The gut virome is thought to play a vital role in shaping and maintaining the composition of the GM and host metabolism^17,18^. It mainly consists of bacteriophages (phages), which are viruses that infect bacteria in a host-specific manner^19^, while archaeal and eukaryotic viruses constitute the remaining. Screening assays can be used to detect known pathogenic viruses before performing FMT. However, in recent years, it has become evident that the human gastrointestinal tract harbors hundreds of eukaryotic viruses with unknown functions^18,20,21^. Although most of these viruses are likely harmless, their potential role in later disease development should not be overlooked, as seen for the human papillomavirus that can induce cervical cancer years after infection^22^.

Independent studies have successfully treated patients suffering from rCDI with sterile filtrated donor feces (containing mainly viruses and limited number of viable bacteria), which were shown to be as effective as FMT^5,23,24^. The successful treatments have been hypothesized to be driven by phage-based restoration of the GM^19,23^. This approach is often referred to as fecal virome transplantation (FVT). In preclinical settings, FVT has also been shown to alleviate symptoms of type-2 diabetes and obesity in male mice^25,26^, prevent the onset of necrotizing enterocolitis in preterm piglets^7^, restore the GM after antibiotic intervention^27^, and improve the proliferation of commensal gut *Akkermansia muciniphila*^28^. These findings highlight the promising application of FVT as a GM restoring treatment targeting various diseases associated with GM dysbiosis. FVT has an advantage over FMT in that it transfers no or very few bacteria and has, compared with FMT, recently been shown to be less invasive for both the gut microbial structure and associated with a lower risk of damaging jejunum in broiler chickens^29^. However, the centrifugation and filtration steps used in preparing FVT cannot separate phages from eukaryotic viruses due to their similar sizes, that together with donor variability, makes widespread use of FVT to treat obesity and type-2 diabetes unlikely. In two recent studies, we aimed to enhance the safety of FVT by developing methodologies that selectively inactivate^30^ (not yet peer-reviewed) or remove the eukaryotic viruses from the fecal matrix^31^ (not yet peer-reviewed) while preserving an active enteric phage community. To achieve this, we utilized the differences in key characteristics between eukaryotic viruses and phages^30,31^; most eukaryotic viruses are enveloped RNA viruses^32,33^ that only infect eukaryotic cells, and most phages are non-enveloped DNA viruses^33,34^ infecting only bacteria. Solvent/detergent treatment (approved by the World Health Organization for treating blood plasma^35^) was used to inactivate enveloped viruses^36^ (FVT-SDT), a compound (pyronin Y) that specifically binds to RNA^37,38^ was applied to inactivate RNA viruses (FVT-PyT), and an optimized chemostat fermentation of intestinal inoculum aimed at removing the eukaryotic viruses by dilution^31^ (FVT-ChP). We have also demonstrated that the modified FVT-SDT and FVT-ChP showed promising results in treating *C. difficile* infections in a mouse model^30^, which represents a simple disease etiology mainly caused by the toxin-producing *C. difficile*^39,40^. The more complex GM associated diet-induced obesity model^41^ was included in this study to investigate whether the same modified FVTs^30^ could be used to improve phenotypes from these two very different disease etiologies through phage-mediated restoration of a dysbiotic GM. Only male mice were included, since female mice are highly protected against diet-induced obesity^42^.

Our main objective was therefore to screen whether the differently modified FVTs^30,31^ had the potential to alleviate symptoms on co-morbidities of metabolic syndrome as a safer and/or more reproducible alternative to the unmodified FVT that we previously have used in a similar diet-induced obesity model^25^.

## Results

We have previously shown that FVT alleviates symptoms associated with metabolic syndrome, including weight gain, impaired glucose tolerance, and MAFLD associated gene expressions in diet-induced obese male mice^25^. As a follow-up study, we hypothesize that different techniques that either deplete or inactivate the eukaryotic viral component in FVT, can improve safety while maintaining the alleviating effects of FVT on symptoms associated with metabolic syndrome^25^. The treatment efficacy of these different modified FVTs (FVT-ChP, FVT-SDT, and FVT-PyT) was compared with unmodified FVT and saline treatment of obese control mice. Lean control mice were also included to evaluate the validity of the conducted diet-induced obesity mouse model. All the transferred intestinal donor content used in the study originated from the same mixed donor material originating from lean male mice. Thus, all results presented only account for male mice and similar results can therefore not necessarily be expected to be observed for female mice.

### Improved clearance of blood glucose and alleviated symptoms of MAFLD following FVT

Oral glucose tolerance tests (OGTT) were conducted on mice after 13 weeks (Fig. S1A-B & Table S1) and 18 weeks (Fig. 1A & 1B) on their respective high-fat or low-fat diet to assess blood glucose regulation. At both study week 13 and week 18, mice on a high-fat diet had significantly (*p* < 0.05) elevated fasting blood glucose levels and impaired glucose regulation during the first 15 minutes after administration compared with the lean control (Fig. 1A & Table S2). Blood glucose levels of mice treated with the chemostat propagated virome (FVT-ChP) decreased sharply from 30 to 60 minutes after glucose administration (Fig. 1A & Table S2), resulting in significantly (*p* = 0.021) improved blood sugar regulation of the FVT-ChP treated mice, compared with the obese control. No FVT treatments improved OGTT scores at the study week 13 (Fig. S1A-B, & Table S1). A MAFLD activity score was determined by liver histopathology (Fig. 1C & S1C-E). Mice treated with unmodified FVT tended to have a reduced (*p* = 0.245) MALFD activity score compared to obese controls, but with a non-uniform response (Fig. S1C-E). Whereas FVT-ChP, FVT-SDT, and FVT-PyT treated mice scored similar to the obese control (Fig. 1C). Weight gain was measured to assess the impact of different treatments on body weight. The obese control mice showed a significant increase (*p* < 0.05) in body weight, MAFLD activity score, epididymal white adipose tissue (eWAT), and blood glucose levels compared to lean controls at the termination of the study (Fig. 1B, 1C, 1D, & 1E), confirming the expected progression of the diet-induced obesity model. Body weight and eWAT size were not improved by any of the FVT treatments compared to obese controls at termination (Fig. 1D and 1E). No mice were excluded in the evaluation of body weight, eWAT size, and liver histopathology. Based on exclusion criteria described in the methods, two mice (FVT-ChP: 1, FVT-SDT: 1) were excluded from the week 13 OGTT measure, and three mice (FVT-ChP: 1, FVT-SDT: 1, unmodified FVT: 1) from the week 18 OGTT measure.

**Fig. 1:**
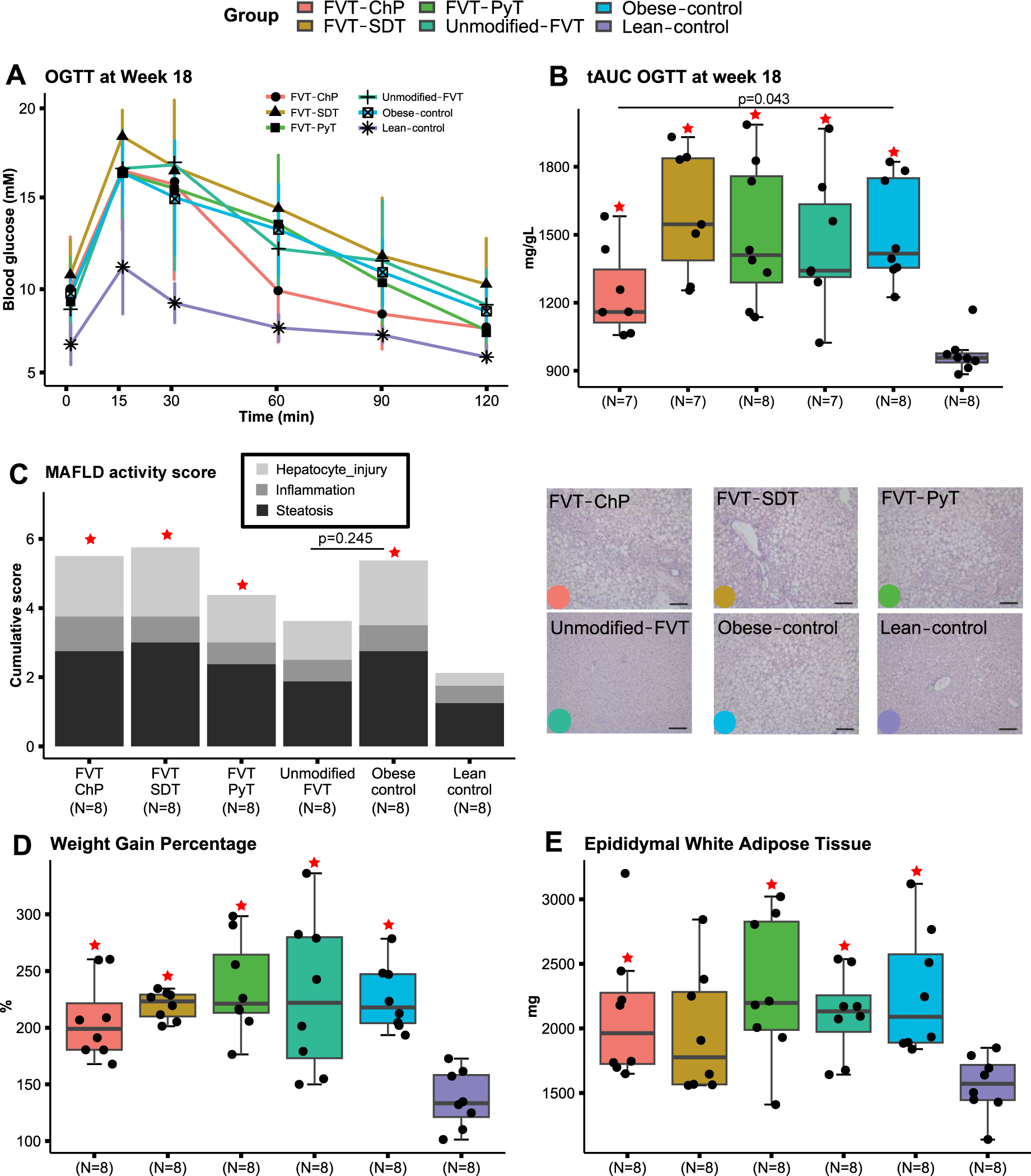
Overview of mouse phenotypic characteristics at termination (23 weeks old) after being treated with modified and unmodified FVT treatments. A) Total area under the curve (tAUC) of the B) OGTT measured at termination of the study, C) MAFLD activity score and representative histology images related to liver tissue pathology. D) Weight gain percentage, and E) measure of eWAT. The labels of *, **, *** between the FVT treatments, obese and lean control mice represent adjusted p < 0.05, 0.01, 0.001 (two-side Wilcoxon rank-sum test with FDR correction), the unadjusted p is presented in figure when it is less than 0.05. Combined boxplots distributions include median, min, max, 25 and 75 percentiles, and outliers (more than 1.5 IQR). Scale in histology images = 300 µm. The * label on top of the boxplot of the treatment represents a p < 0.05 between the treatment and the lean control mice (two-side Wilcoxon rank-sum test).

### Unmodified fecal virome affected the immune response in adipose tissue

Fluorescence-activated cell sorting (FACS) was performed to evaluate the proportions of immune cells in eWAT (Fig. 2A-L) at termination (study week 18). Interestingly, mice treated with the unmodified FVT showed a decrease in the proportions of dendritic cells (%CD11c+ of CD45+) (*p* = 0.05), T helper cells (%CD4+ of TCRab+) (*p* = 0.015), activated macrophages (%CD11c+ of F4.80+) (*p* = 0.038), and central memory CD8+ T cells (%CD44+CD62l+ of CD8+) (*p* = 0.05) compared to the obese control (Fig. 2A, 2C, 2E & 2G). Except for mice treated with unmodified FVT, mice fed a high-fat diet generally appeared with a significant (*p* < 0.05) increase in the proportions of dendritic cells (% CD11c+ of all CD45+ leukocytes), central memory CD8+ T cells (%CD44+CD62l+ of CD8+), naïve CD8+ T cells (%CD44-CD62l+ of CD8+), and in the proportion of activated macrophages (%CD11c+ of F4.80+ macrophages), compared with the lean control group (Fig. 2A, 2G, 2I & 2C). The proportions of immune cells in the mesenteric lymph node (MLN) were also measured but showed no clear differences between treatment groups (Fig. S2). FACS analysis failure resulted in the exclusion of three mice (FVT-ChP: 1, FVT-SDT: 1, lean control: 1) from the FACS analysis of adipose tissue and six mice from MLN tissue (lean control: 2, obese control: 1 FVT-PyT: 1, unmodified FVT: 2).

**Fig. 2:**
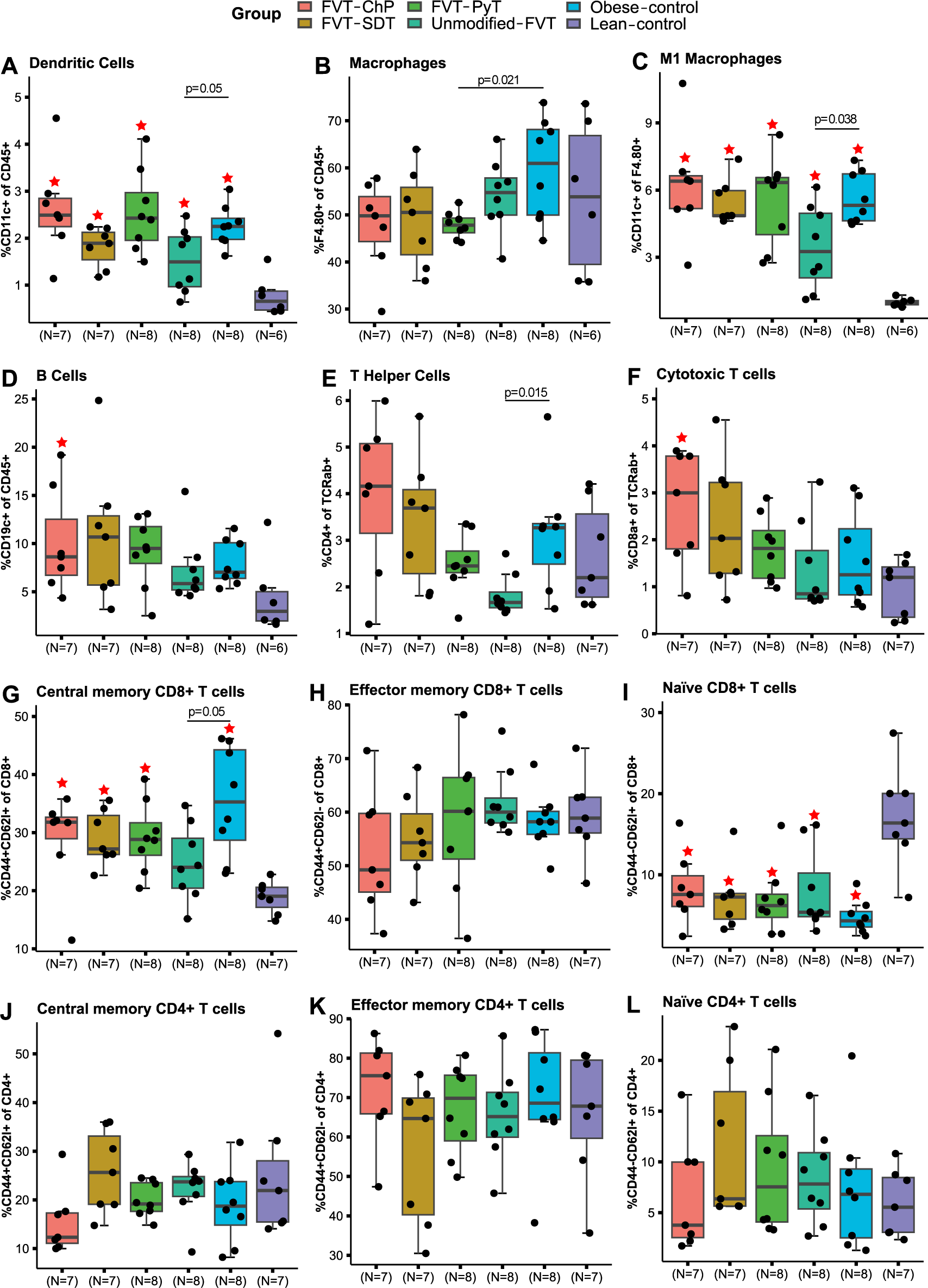
Proportions of immune cells in adipose tissue at study termination (23 weeks old). A) to L) are showing the overall FACS profile in the mouse adipose tissue. The labels of *, **, *** between the FVT treatments, obese and lean control mice represent adjusted p < 0.05, 0.01, 0.001 (two-side Wilcoxon rank-sum test with FDR correction), the unadjusted p is presented in figure when it is less than 0.05. Combined boxplots distributions include median, min, max, 25 and 75 percentiles, and outliers (more than 1.5 IQR). The * label on top of the boxplot of the treatment represents a p < 0.05 between the treatment and the lean control mice (two-side Wilcoxon rank-sum test).

### Inflammatory signal associated with the chemostat-propagated virome

Motivated by the observed effects on OGTT and histological measures by the unmodified FVT and FVT-ChP treated mice compared with the lean and obese control groups, we decided only to examine the cytokine profile of these four groups (representing a total of 32 mice). The serum cytokine profile of FVT-ChP treated mice showed significantly elevated (*p* < 0.05) expression of three pro-inflammatory cytokines, including IL-15, TNF-α, and MIP-2 compared to the obese control (Fig. S3). The overall cytokine profile of serum from mice treated with unmodified FVT was not different from that of obese control mice. Therefore, the cytokine profile does not explain the improved liver pathology (Fig. 1C). It should be noted that interpretation of the cytokine profiling is challenged by the lack of significant differences between the obese and lean control in 9 out of 10 investigated cytokines.

### Shifted gut bacteriome composition was linked to metabolic syndrome improvement

Upon arrival to our housing facility, the bacterial composition of the mice appeared comparable based on bacterial diversity (Shannon diversity index) and bacterial composition (Bray-Curtis dissimilarity) (Fig. 3A & 3B). The change in diet and environmental conditions in our housing facilities, compared with that of the vendor, clearly affected the bacterial diversity (Fig. 3A) of all groups, including the lean control. However, the bacterial diversity for the lean control was close to normalized at termination compared with the arrival. As expected, the *ad libitum* low-fat diet provided to the lean control mice, significantly (*p* < 0.05) affected the bacterial diversity and composition compared to the *ad libitum* high-fat diet fed groups (Fig. 3A & 3B). At the termination of the study, the FVT-ChP treated mice had a significantly lower (*p* = 0.028) bacterial diversity compared with obese control mice (Fig. 4A & 3A). Except for FVT-SDT (*p* = 0.12), all the different FVT-treated mice harbored a significantly (*p* < 0.05) altered gut bacterial composition compared to the obese control group at termination (Fig. 3B). The dominant bacterial phyla detected in the mice feces were *Firmicutes*, *Bacteroidetes*, *Proteobacteria*, *Verrucomicrobia*, *Deferribacteres*, and *Actinobacteria* (Fig. S4J). We adopted DESeq2 to identify differentially abundant bacterial taxa between the FVT treatment groups and obese control mice. The FVT-ChP treated mice were observed with a significant (*p* < 0.05) increase in the relative abundance of *Limosilactobacillus reuteri*, and a tendency (*p* < 0.1) of increase in bacterial taxa belonging to lactobacilli, *Allobaculum*, and *Bacteroidales* compared with the obese control mice (Fig. 3D). In contrast, unmodified FVT treated mice had a significant (*p* < 0.05) decrease in the relative abundance of bacteria belonging to the genus *Allobaculum* and a tendency (*p* < 0.1) of increase was observed for *Erysipelotrichaceae* and *Bacteroidales* (Fig. 3D).

**Fig. 3:**
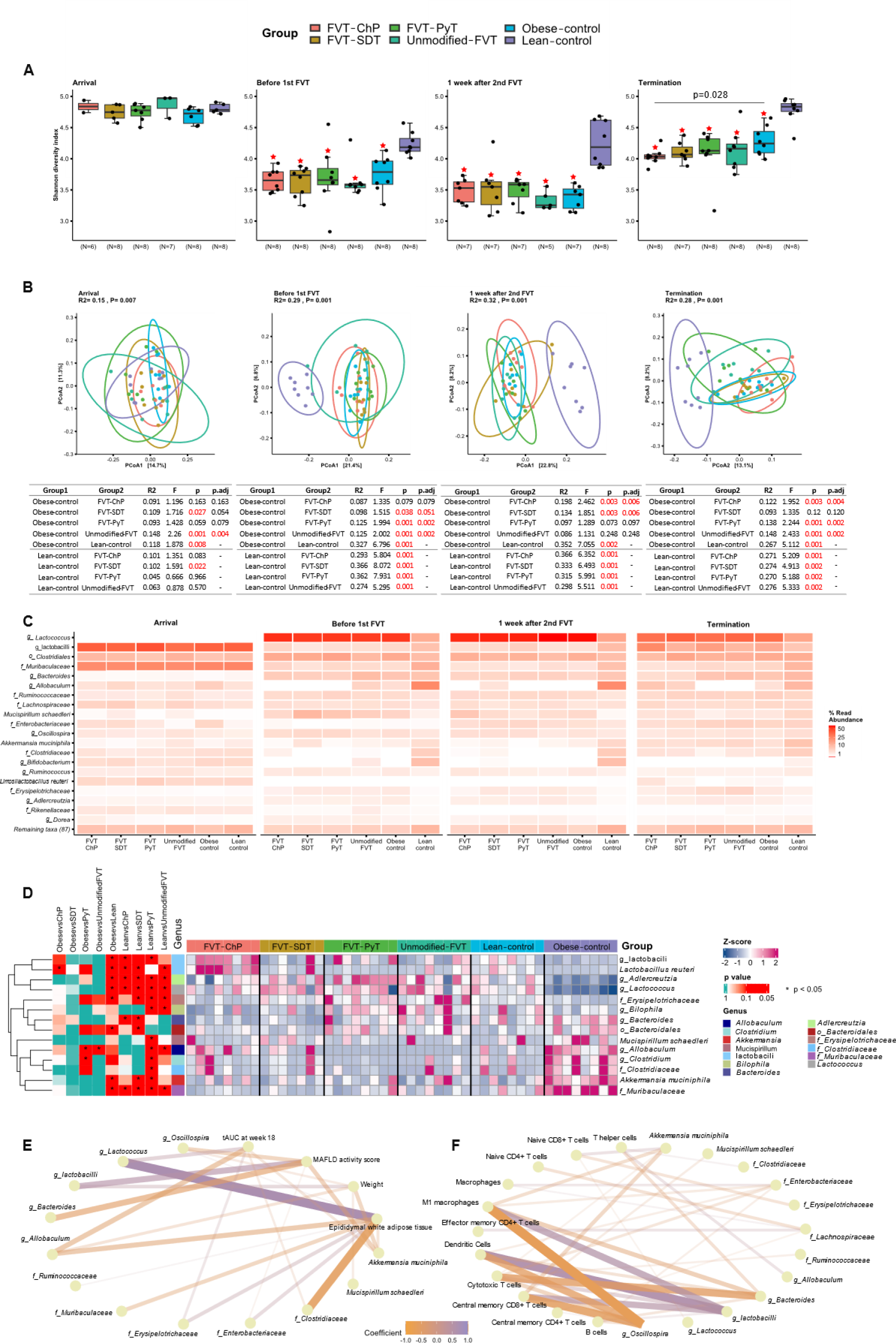
Bacteriome analysis based on 16S rRNA gene amplicon sequencing at four time points: at arrival (before diet intervention (5 weeks old), before 1^st^ FVT (11 weeks old), one week after 2^nd^ FVT (14 weeks old), and study termination (23 weeks old). A) The bacterial diversity (Shannon diversity index), combined boxplots distributions include median, min, max, 25 and 75 percentiles, and outliers (more than 1.5 IQR), B) PCoA plot of the bacterial composition (Bray-Curtis dissimilarity), alongside a table showing pairwise PERMANOVA results (*p* adjusted for FVT treatments and obese control mice comparisons by FDR correction), C) heatmap representing the relative abundance of dominating bacterial taxa, D) heatmap highlighting significant (*p* < 0.05) difference in the differentially abundant bacterial taxa, and E) to F) Pairwise Spearman’s correlations between relative abundance of bacterial taxa (relative abundance > 1%) and phenotypic characteristics, and immune cell proportions in adipose tissue, after FDR correction for multiple comparison. The absolute values of coefficients are shown in the figure. The zOTUs were collapsed into the best possible taxonomical resolution. The labels of *, **, *** between the FVT treatments, obese and lean control mice represent adjusted p < 0.05, 0.01, 0.001 (two-side Wilcoxon rank-sum test with FDR correction), the unadjusted p is presented in figure when it is less than 0.05. Combined boxplots distributions include median, min, max, 25 and 75 percentiles, and outliers (more than 1.5 IQR). The * label on top of the boxplot of the treatment represents a p < 0.05 between the treatment and the lean control mice (two-side Wilcoxon rank-sum test).

**Fig. 4:**
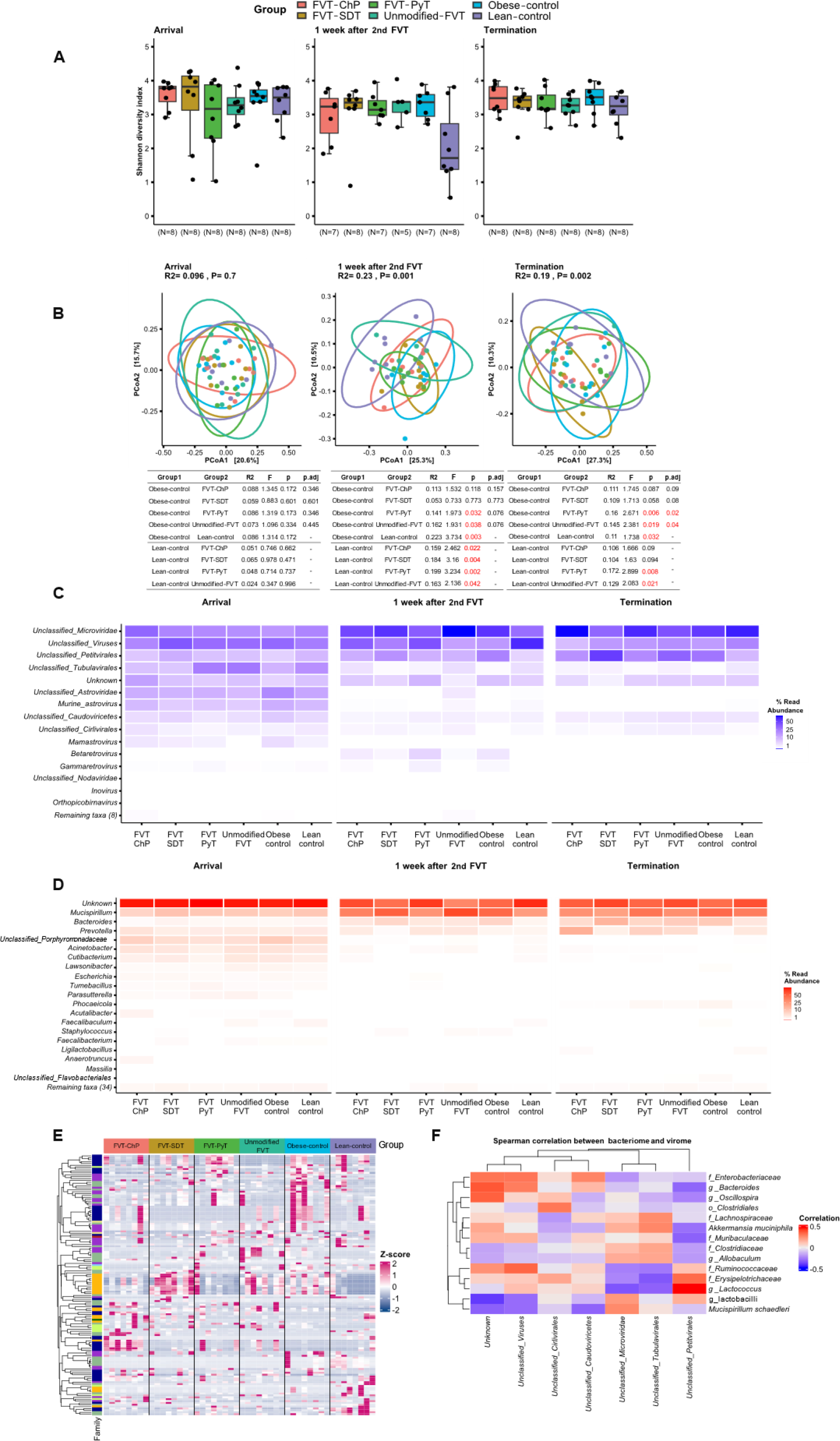
Metavirome analysis based on whole-genome sequencing at three time points: at arrival (before diet intervention (5 weeks old), one week after 2^nd^ FVT (14 weeks old), and study termination (23 weeks old). A) Boxplots showing the viral diversity (Shannon diversity index), B) PCoA plot of the viral composition (Bray-Curtis dissimilarity), alongside a table showing pairwise PERMANOVA results (*p* adjusted for FVT treatments and obese control mice comparisons by FDR correction), C) heatmap representing the relative abundance of dominating viral taxa, D) heatmap representing the relative abundance of the predicted bacterial hosts (based on the viral contigs), E) heatmap highlighting significant (*p* < 0.05, FDR correction) difference in the differentially abundant viral contigs based on vOTUs level, F) Pairwise Spearman’s correlations between relative abundance of bacterial species (relative abundance > 1%) and viral genus (relative abundance > 0.1%), with the FDR correction for multi-comparison. The viral contigs were collapsed into the best possible taxonomical resolution. The labels of *, **, *** between the FVT treatments, obese and lean control mice represent adjusted p < 0.05, 0.01, 0.001 (two-side Wilcoxon rank-sum test with FDR correction), the unadjusted p is presented in figure when it is less than 0.05. Combined boxplots distributions include median, min, max, 25 and 75 percentiles, and outliers (more than 1.5 IQR). The * label on top of the boxplot of the treatment represents a p < 0.05 between the treatment and the lean control mice (two-side Wilcoxon rank-sum test).

### Altered bacterial compositions were associated with improvements of metabolic syndrome

Correlation analyses were performed to elucidate potential links between the bacterial GM component and the alleviating effects associated with the unmodified FVT and FVT-ChP. Using phenotypic characteristics and measured proportions of immune cells in adipose tissue as indicators of metabolic syndrome of the mice, we performed Spearman correlation analysis (FDR corrected p < 0.05, |Spearman coefficient| > 0.4) between these indicators and the most abundant bacterial taxa (relative abundance > 1%). The improved glucose regulation was positively correlated with the relative abundance of *Allobaculum spp.* (coefficient = -0.47) and negatively correlated with the relative abundance of *Lactococcus spp.* (coefficient = 0.41) (Fig. 3E). The elevated relative abundance of *A. muciniphila* (coefficient = -0.43), *Bacteroides spp*. (coefficient = -0.51), *Oscillospira spp.* (coefficient = -0.42) were correlated with lower MAFLD activity scores. In contrast, the relative abundance of unclassified lactobacilli was positively correlated (coefficient = 0.52) with the MAFLD activity score (Fig. 3E). When the size of eWAT increase, the relative abundance of taxa *Allobaculum spp.* (coefficient = -0.46) tended to be diminished, while the relative abundance of *Lactococcus spp.* (coefficient = 0.68) decrease (Fig. 3E). The bodyweight gain of the mice was negatively correlated (coefficient = - 0.41) with the relative abundance of *A. muciniphila* (Fig. 3E).

The pairwise Spearman correlations were established between the proportions of immune cells from adipose tissue and bacterial taxa. The abundance of unclassified lactobacilli exhibited a positive correlation with the proportions of activated macrophages, central memory CD8+ T cells, dendritic cells, and cytotoxic T cells (respective coefficients = 0.54, 0.52, 0.63, and 0.41) (Fig. 3F). Conversely, the abundance of *Oscillospira spp.* demonstrated an inverse correlation with the proportions of four immune cells (respective coefficients = -0.65, -0.59, -0.56, and - 0.42) (Fig. 3F). The negative correlation was also found between *Bacteroides spp.* and proportions of activated macrophages (coefficient = -0.44), as well as cytotoxic T cells (coefficient = -0.57) (Fig. 3F).

### Transplantation of modified fecal viromes shifted the GM composition

None of the FVT treatments affected gut viral diversity (Shannon diversity index), and neither any differences between the lean and obese control (Fig. 4A). The viral composition (Bray-Curtis dissimilarity) of mice treated with the different FVTs was different (*p* < 0.1) from the obese control at termination (Fig. 4B). This indicated that introducing a new viral community through FVT leads to changes in the recipient’s gut viral composition. The taxonomic profiles of the recipient gut viromes were dominated by unclassified viruses belonging to the family *Microviridae*, order *Petitvirales* and order *Tubulavirales*-associated viruses (Fig. 4C), and the predicted (based on viral contigs) bacterial hosts were dominated by the genera of *Mucispirillum*, *Bacteroides*, and *Prevotella* (Fig. 4D). Differential viral relative abundance was analyzed at the level of viral contigs (vOTUs) (Fig. 4E & S5) to support the observed differences in the viral composition. When comparing both FVT-ChP and unmodified FVT treated mice to the obese control at the study termination, significant (*p* < 0.05) differences were observed in the relative abundance of viruses belonging to the family *Microviridae*, order *Petitvirales,* and class *Caudoviricetes* (Fig. 4E & S5). Pairwise Spearman’s correlation analysis was conducted to investigate the influence of viral composition on the gut bacteriome. The relative abundance of different viral contigs of the family *Microviridae* both negatively (coefficient < -0.4) correlated with the genus *Allobaculum* and positively (coefficient > 0.4) correlated with lactobacilli, *Bacteroides spp.*, and the family *Clostridiaceae* (Fig. 4F & S6). Furthermore, the viral contigs of order *Petitvirales* showed a strong positive (coefficient > 0.4) correlation with the genus of *Lactococcus* (Fig. 4F & S6). Eukaryotic viruses appeared to be nearly depleted in the FVT-ChP in terms of relative abundance (0.1%) compared to the FVT-SDT (3.66%), unmodified FVT (1.07%), and FVT-PyT (7.41%) (Fig. S10E). It should be emphasized that the metavirome sequencing can solely be applied to evaluate the removal of viruses, while it cannot differentiate whether viral particles have been inactivated or not from e.g. the solvent-detergent or pyronin Y treatment, and the relative abundance does not account for the quantity of viruses. Fecal samples of mice were excluded from bacteriome (11 samples) and virome (6 samples) analysis due to lack of sample material or low sequencing reads (Fig. S12).

### Recipient-dependent response to unmodified FVT

In this study, mice treated with unmodified FVT revealed a tendency of a reduction in the MAFLD activity score compared with obese control mice (Fig. 1C). However, this reduction was not uniformly observed across all mice within the unmodified FVT treated mice (Fig. S1C– E). Instead, it was predominantly driven by a subset of mice exhibiting a more pronounced cage independent response to the unmodified FVT treatment (Fig. S1C, S1D & S1E). To understand what caused the different responses to the same treatment, the study subjects were stratified into two distinct groups: those exhibiting a response to the unmodified FVT treatment (unmodified-FVT response; characterized by hepatocyte injury score ≤ 1, inflammation score ≤ 1, and steatosis ≤ 2; number of mice = 4), and those showing no response (unmodified-FVT non-response; with hepatocyte injury score > 1, inflammation score > 1, and steatosis > 2; number of mice = 4).

Despite the low sample size, mice within the unmodified-FVT responder group displayed a significant (*p* < 0.05) downregulation in MAFLD severity (Fig. 5A) and a tendency of improved blood glucose regulation relative to the obese controls (Fig. 5D). The unmodified-FVT response group also demonstrated significantly (*p* < 0.05) reduced proportions of dendritic cells (%CD11c+ of CD45+ leukocytes), activated macrophages (%CD11c+ of F4.80+), T helper cells (%CD4+ of TCRab+), and central memory CD8+ T cells (%CD44+CD62l+ of CD8+) in the adipose tissue, compared to the obese control mice (Fig. S7A, S7C, S7E, & S7G). No clear differences were observed in the proportions of immune cells in the mesenteric lymph node (Fig. S8) or the blood serum cytokine profile (Fig. S9). Moreover, the bacterial profile of the mice responding to the unmodified FVT treatment was significantly separated from the non-responding (*p* = 0.028) and the obese control mice (*p* = 0.005) (Fig. 5F). This divergence was further underscored by the increased relative abundance of *Bacteroides* and *Dehalobacterium*, alongside a significant (*p* < 0.05) increase in *Oscillospira* and a decrease in lactobacilli in the unmodified-FVT responder group relative to the obese controls (Fig. 5H, 5K, 5L, and 5N). These findings imply that responses to the same treatment regimen can vary due to individual differences, however, further studies need to be conducted to validify these observations, since the animal model was neither designed nor hypothesized to include the responder versus non-responder stratification.

**Fig. 5:**
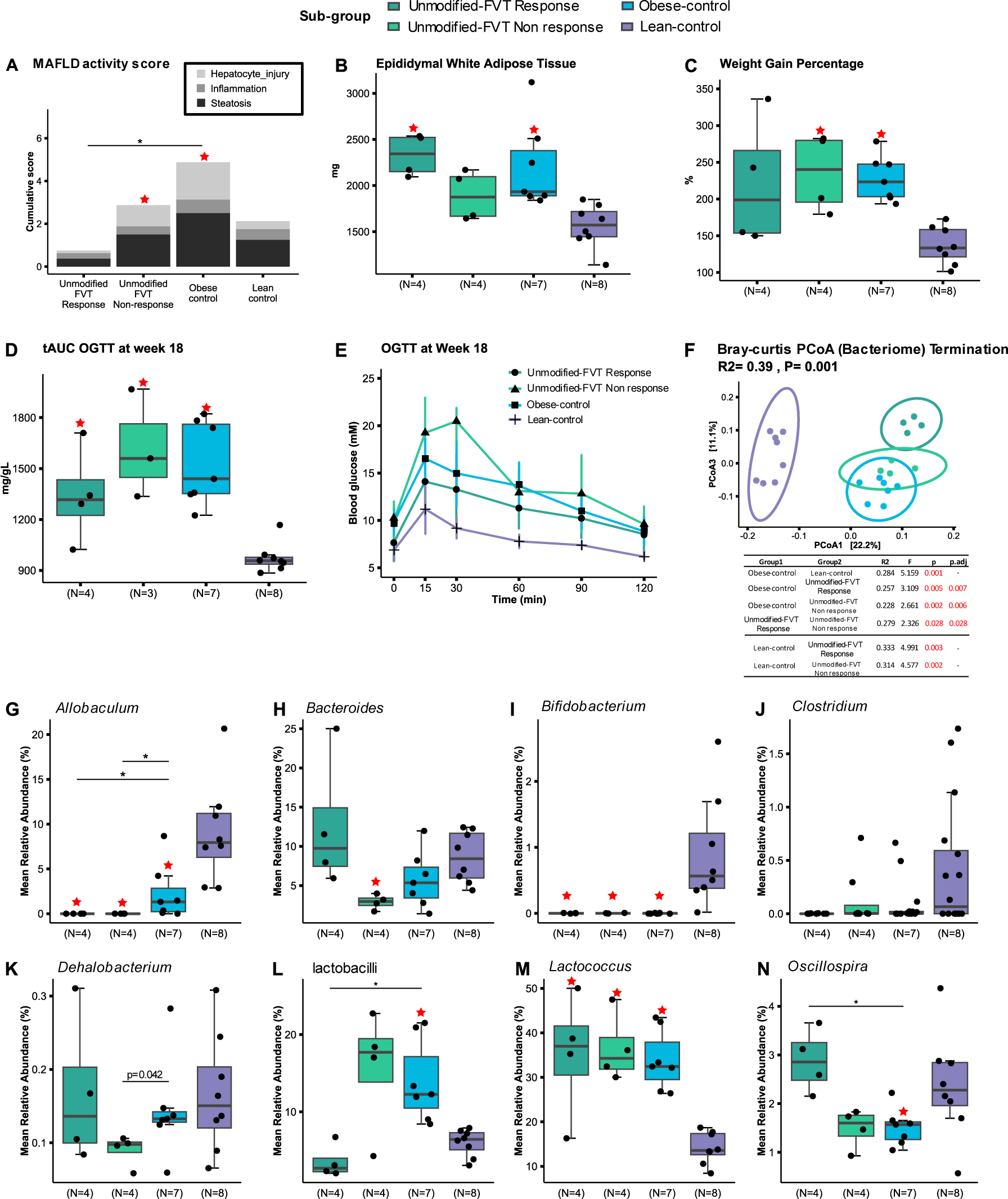
Overview of the unmodified FVT treatment stratification analysis based on response and non-response groups at termination (23 weeks old). A) MAFLD activity score and representative histology images related to liver tissue pathology, B) weight gain percentage, C) measure of eWAT, D) Total area under the curve (tAUC) of the E) OGTT measured at termination of the study, F) PCoA plot of the bacterial composition (Bray-Curtis dissimilarity), alongside a table showing pairwise PERMANOVA results (*p* adjusted for FVT treatments and obese control mice comparisons by FDR correction), and G) to N) boxplots showing the relative abundance of selected bacterial taxa. The labels of *, **, *** between the FVT treatments, obese and lean control mice represent adjusted p < 0.05, 0.01, 0.001 (two-side Wilcoxon rank-sum test with FDR correction), the unadjusted p is presented in figure when it is less than 0.05. Combined boxplots distributions include median, min, max, 25 and 75 percentiles, and outliers (more than 1.5 IQR). The * label on top of the boxplot of the treatment represents a p < 0.05 between the treatment and the lean control mice (two-side Wilcoxon rank-sum test).

## Discussion

To improve the safety of fecal virome transplantation (FVT), recently developed modification methodologies were applied to inactivate or remove the eukaryotic viral component of FVT while maintaining an active enteric phage community^30,31^. We examined the potential of three different modified FVTs to mitigate symptoms of metabolic syndrome compared with unmodified FVT and saline, using a diet-induced obesity male mouse model. Mice treated with a chemostat-propagated virome (FVT-ChP) that was nearly depleted from eukaryotic viruses by dilution, expressed a significantly (*p* < 0.05) improved blood glucose regulation compared with the obese control. Considering the reproducibility potential of chemostat propagated enteric phageomes^31,43^, refinement of this FVT modification strategy may open interesting perspectives for enhancing the reproducibility of FVT-based studies and safety by significantly reducing the number of eukaryotic viruses transferred. We have previously shown that unmodified FVT could reduce weight gain and normalize blood glucose regulation in male mice^25^, but this was only partly replicated in the present study with a non-uniformly cage-independent phenotypical response of mice treated with unmodified FVT. Mice that responded on the unmodified FVT treatment were characterized by improved oral glucose tolerance test (OGTT) measures, a low liver histopathology, and the bacterial composition was distinctively separated from the non-responding mice and the obese control. This may be in line with our earlier observations where unmodified FVT showed capable to affect the expression of genes with important metabolic liver functions to be more like the lean control than the obese control mice^25^. The observed discrepancies between the previous^25^ and the present study are likely attributable to variations in the gut microbiome (GM) of the donor^44^. Such variations can differ across mouse vendor facilities, posing challenges to reproducibility^45–48^. Additionally, while differences in the GM of the recipient contribute to these discrepancies, their impact might be less pronounced compared to the variability originating from donor materials^49,50^.

The cytokine profile of the blood serum was examined since low-grade systemic inflammation is connected to metabolic syndrome^51,52^. Especially elevated levels of the cytokines MIP-2, IL-15, and TNF-α in the FVT-ChP treated mice indicated elevated inflammation. However, the interpretation was challenged by inconsistent differences in the cytokine levels between the lean and obese control groups (Fig. S3). The recruitment and activation of neutrophils via MIP-2 can prompt the release of diverse inflammatory mediators, which can hasten the onset of liver inflammation^53^. TNF-α is produced by adipose tissue and works as a pro-inflammatory cytokine that has been shown to play a role in the development of metabolic dysfunction-associated fatty liver disease (MAFLD)^54,55^. Overexpression of IL-15 in transgenic mice and IL-15 treatment of NOD mice have been reported to improve glucose tolerance^56^. As a double edge sword, the increased levels of pro-inflammatory cytokines (MIP-2, IL-15, and TNF-α) in the blood serum of FVT-ChP treated mice compared to the obese control may partly explain their improved blood glucose regulation, as well as the lack of improvement in their associated weight gain measures and histopathology liver score compared to the obese control mice.

Phage immunogenicity has garnered increased interest due to its potential role in regulating our immune system by sharing similar structures and proteins with eukaryotic viruses^57^. Phages may be both passively and actively distributed throughout various parts of our body^58^. The interactions between phages transferred along with the chemostat-propagated virome and the host immune system could help explain our observations of increased levels of three out of ten cytokines in the FVT-ChP treated mice, compared with the obese control. However, we lack evidence of specific viruses that could have triggered this immune response in the blood serum. Although low-grade systemic inflammation is connected to metabolic syndrom^51,52^, it does not exclude the possibility of improved glucose regulation alongside increased levels of some cytokines in blood serum.

The inflammatory state of obesity is associated with an accumulation of macrophages in the adipose tissue^59^, where the activated macrophages are recruited by CD8^+^ T cells^60^. Additionally, elevated levels of dendritic cells can play a role in the immune response to liver injury^61^. In this study, reductions in the proportions of CD8+ T cells, activated macrophages, T helper cells, and dendritic cells were observed in the mice treated with unmodified FVT compared to the obese control, which may be linked to the observed reduction in the liver pathology. Since we do not have the total cell count from adipose tissue and there were no significant differences in cytokine levels (Fig. S3 & S9), it is challenging to draw a general conclusion about the extent to which there was less inflammation in the unmodified FVT group. However, the phenotypical differences present in the liver tissue, adipose tissue, and blood cytokine profile all point in the same direction.

It is commonly accepted that high-fat diet^62^ and housing conditions^45,47,63^ substantially affects the GM of mice, which would explain the sudden overall decrease in the bacterial diversity in all groups both before and after the study intervention. The relative abundance of *Allobaculum spp.* was increased (*p* = 0.083) in feces samples from FVT-ChP treated mice and significantly (*p* < 0.05) decreased in unmodified FVT treated mice compared with the obese control (Fig. S4A). *Allobaculum* is a Gram-positive bacteria found in both murine and human hosts^64,65^. Prior studies have consistently identified impaired intestinal integrity, thickness of mucus layer, and immunity (low-grade inflammation) as important factors contributing to the development of metabolic syndromes related diseases^66–69^. Mucin degrading bacteria like *Akkermansia muciniphila*^67,70,71^ and *Allobaculum* spp.^72–74^ have been reported to play a key role in these processes. *Allobaculum* has been reported as a particularly active glucose utilizer that produces lactate, butyrate^75^, and impacts the metabolism of long chain fatty acids^76^. Through the production of short-chain fatty acids like butyrate, *Allobaculum* contributes to the protection of the intestinal barrier^72^ stimulates the immune system, and putatively protects against metabolic syndrome^73^. This would be in accordance with supplementation of butyrate limiting hyperglycemia through the regulation of amongst glucagon-like peptide-1 (GLP-1) and insulin in serum^77^. This is further supported by *Allobaculum* being reported to be positive correlated to hypoglycemia and negative correlated to HOMA-IR levels^78^ (not yet peer-reviewed). Taken together, the current understanding of the role of *Allobaculum* on metabolic syndrome supports our observations of improved glucose regulation for the FVT-ChP treated mice that had increased relative abundance of *Allobaculum* compared with obese control (Fig. S4A).

Homologs of four enzymes involved in the butyrate pathway have been discovered in the genome of species belonging to genus of *Oscillospira* suggesting it as a potential contributor to butyrate production^79^. Thus, the significant increase (*p* < 0.05) in the relative abundance of *Oscillospira* (Fig. 5G) in the mice that responded on the unmodified FVT compared with the obese control, may have played an important role in the butyrate production in these animals. *Oscillospira* could be speculated to have a redundant metabolic role in the butyrate production along with *Allobaculum,* since *Allobaculum* was not detectable in the unmodified FVT treated mice (Fig. 5N). This might have contributed to the tendency of an enhanced blood glucose clearance in the mice that responded on the unmodified FVT (Fig. 5D & 5E). The level of *Oscillospira* spp. has also been negatively associated with hepatic fat accumulation in pediatric cases of MAFLD^80^ and non-alcoholic steatohepatitis (NASH)^81^, hence indicating a protective role by *Oscillospira* of fat accumulation in the liver.

The determination of the differential viral abundance showed clear differences when comparing FVT-ChP treatment with unmodified FVT (Fig. S5). Thus, these differences in viral profiles may also have contributed to the development of the two different observed phenotypes. The mechanisms behind the GM modulating effects of FVT are still poorly understood, but accumulating reports suggest that the phenotypic traits of FVT donors, to some extent, can be transferred to recipients, as the phages might catalyze a modulation of the recipient ecosystem to similar to their origin. This is exemplified by phage donor profiles being transferred to the gut of *C. difficile* patients after successful FMT/FVT^23,82,83^, FVT from lean mice could shift the GM composition in obese mice to resemble that of lean individuals^25^, and FVT donor material originating from an ecosystem with a relatively high abundance of *A. muciniphila*, could significantly increase the abundance of the enteric endogenous (native) *A. muciniphila* in mice that received the FVT^28^. This may be driven by cascading events^19^, as demonstrated in a gnotobiotic mouse model^84^, where phage infections indirectly influence the bacterial balance. Thus, the far more complex viral composition of FVT could similarly alter the bacterial ecosystem and as observed in our previous study, affect the blood metabolome and GM, leading to systemic effects^25^. This concept might seem counterintuitive given the general belief in the strain-specific nature of phages, however, a recent study proposed that phages could interact with distantly related microbial hosts^85^. Phage satellites have, amongst other, been suggested to contribute to broader host ranges^86,87^. Also the transfer of potentially beneficial metabolic genes from temperate phages to their bacterial hosts^88–91^, may enhance host competitiveness and contribute to overall GM changes. These findings align with recent research demonstrating how metabolic functions and bacterial interaction networks were context-dependent of variables like nutrition, host environment, and bacterial compositions^92^, supporting the hypothesis that cascading events initiated by FVT could catalyze GM modulating effects. In addition to the bacteria-phage relations, the impact of the immune system on gut health should not be neglected. Recent evidence suggests that bacteriophages interact with our immune system^57,58^ through mechanisms like TLR3 and TLR9^93,94^. Stimulation of the immune system may thus be another mechanism behind the efficacy of FVT.

Both the modified FVT-SDT (3.66%) and FVT-PyT (7.41%) exhibited a notably higher relative abundance of eukaryotic viruses when compared to the unmodified FVT (1.07%). The inactivation process involving solvent detergent and pyronin Y might also lead to the destruction of a certain fraction of phages. Consequently, this could impact the relative abundance distribution between phages and eukaryotic viruses. While further experimental data is required to substantiate this hypothesis, it aligns with our other study that delves into the efficacy of these methods^30^. In this parallel investigation, pyronin Y reduced the phage activity of the majority of the examined phages, whereas solvent/detergent only affected the activity of one of the tested phages. The chemostat fermentation setup aimed to mimic mouse gut conditions^31^ and utilized the same intestinal content as other FVT treatments. However, the chemostat propagation will inevitably alter phage composition and diversity compared to the original inoculum, as also illustrated by the differences in phage composition in the chemostat-propagated virome compared to other treatments (Fig. S10), also offering a possible explanation of the decreased Shannon diversity index observed in the FVT-ChP group, compared to the obese control. The chosen sample size (n = 8) group housed in two cages, is previously validated as sufficient for diet-induced obesity^95^, thereby accommodating the 3R principle of reducing the number of animals^96^. The combination of group housing and the coprophagic behaviour of the mice may cause in cage-associated effects^97^, which constitutes a limitation of the study. However, statistical analysis of the phenotypical measures (Table S4) showed no pronounced cage-associated effects concerning the main findings. A recent report suggests to decrease animal density in the cages to increase the statistical power^97^, which could address the cage-associated variance issue while maintaining accommodation of the 3Rs^96^ in future studies. The viral diversity across all groups showed no cage effects for the Shannon diversity index. Similarly, the viral composition in all FVT groups remained unaffected by cage effects, while the lean and obese control groups displayed cage-associated effects. Regarding bacteria, only the Shannon diversity index of the lean control group showed cage-associated effects. However, bacterial composition in most groups exhibited cage-associated effects, likely influenced by coprophagic behavior, a common issue in mouse studies^97,98^. Thus, it cannot be ruled out that cage-associated differences in the bacterial composition may have influenced the treatment efficacy of the different FVTs.

Most drug-based treatments for chronic diseases are long-term or periodic^99^, but this study only treated mice with FVT twice. Compared with the phenotypical changes induced by a long-term high-fat diet, two FVT treatments may be insufficient, raising the question of whether the number and frequency of FVT treatments should be adjusted in future intervention studies.

To summarize, it was possible to partly maintain the alleviating effects of FVT^25^ on metabolic syndrome-associated symptoms with a reproducible chemostat-propagated enteric virome that was nearly depleted from eukaryotic viruses. While the unmodified FVT showed the most substantial phenotypic change by reducing diet-induced liver tissue damage and lowered the proportions of immune cells in the adipose tissue. The mice that received the unmodified FVT either responded to the treatment or did not. In contrast, the two other FVT modifications (FVT-SDT and FVT-PyT) did not appear as promising modification strategies of FVT targeting diet-induced obesity, since no clear phenotypical improvements were observed.

The concept of chemostat-propagated phageomes (FVT-ChP) could address the main limitations that are associated with unmodified FVT: procuring and reproducing sufficient enteric phage solutions, *in vitro* donor-recipient compatibility screening of the individual GM, and minimizing the transfer of eukaryotic viruses from donors to recipients. Considering that the concept of FVT-ChP as a modification of FVT already in this premature stage has demonstrated phenotypical effects in treating two distinct disease etiologies of diet-induced obesity and *C. difficile* infection^30^, it urges for refinements as a therapeutic tool targeting other diseases associated with GM dysbiosis.

## Methods

### The animal origin and preparation of donor viromes

A total of 54 male C57BL/6N mice were purchased for the purpose of harvesting intestinal content for downstream FVT applications. Upon arrival, the mice were five weeks old and obtained from three vendors: 18 C57BL/6NTac mice from Taconic (Denmark), 18 C57BL/6NRj mice from Janvier (France), and 18 C57BL/6NCrl mice from Charles River (Germany). The mice were earmarked upon arrival, randomly (simple randomization) assigned according to vendor to 3 cages with 6 mice each, and housed at the Section of Experimental Animal Models, University of Copenhagen, Denmark, following previously described conditions^48^. They were provided with *ad libitum* access to a low-fat diet (LF, Research Diets D12450J) for a period of 13 weeks until they reached 18 weeks of age, which was the planned termination point. Unfortunately, malocclusions resulted in malnutrition for two C57BL/6NRj mice, and they were euthanized before the intended termination date. All mice were euthanized by cervical dislocation, and samples of intestinal content from the cecum and colon were collected and suspended in 500µL of autoclaved anoxic PBS-buffer (NaCl 137mM, KCl 2.7mM, Na_2_HPO_4_ 10mM, KH_2_PO_4_ 1.8mM). Subsequently, all samples were stored at -80°C. In order to preserve the viability of strict anaerobic bacteria, 6 mice from each vendor (a total of 18 mice) were sacrificed and immediately transferred to an anaerobic chamber (Coy Laboratory) containing an atmosphere of approximately 93% N_2_, 2% H_2_, and 5% CO_2_, maintained at room temperature. The samples collected from these mice within the anaerobic chamber were used for anaerobic chemostat cultivation to produce the chemostat propagated virome (FVT-ChP). The intestinal content from the remaining 34 mice was sampled under aerobic conditions and used to generate the fecal virome for downstream processing of the unmodified FVT, FVT-SDT, and FVT-PyT treatments. A flow diagram illustrating the aforementioned processes is provided (Fig. S11). All procedures for handling animals used for collecting donor material were carried out in accordance with the Directive 2010/63/EU and the Danish Animal Experimentation Act with the license ID: 2012-15-2934-00256.

### Unmodified fecal virome

Intestinal content from cecum and colon was thawed and processed to produce FVT solutions as previously described^25^, with the exception of Centriprep Ultracel YM-30K units (Millipore) being replaced with YM-50K units, which is applied to concentrate fecal viromes and remove metabolites below the size of 30 kDa. Fecal viromes of mice from all vendors were mixed and divided into three aliquots. One was immediately stored at -80°C and represented the unmodified FVT. The two-remaining fecal virome aliquots were further processed for inactivation of eukaryotic viruses by either dissolving the lipid membrane of enveloped viruses with solvent/detergent treatment or inhibiting replication of RNA viruses with pyronin Y treatment.

### Solvent/detergent-treated fecal virome (FVT-SDT)

The solvent/detergent treatment is commonly used for inactivating enveloped viruses (most eukaryotic viruses are enveloped) in blood plasma, while non-enveloped viruses (most phages are non-enveloped) are not inactivated^36,100^. The fecal viromes were treated according to the recommendations from the WHO for clinical use of solvent/detergent treated plasma; incubation in 1% (w/v) tri(n-butyl) phosphate (TnBP) and 1% (w/v) Triton X-100 at 30°C for 4 hours^35^. Solvent/detergent treatment was performed by following the method of Horowitz *et al.*^36^ and the removal of TnBP and Triton X-100 was performed as described by Treščec *et al.*^101^. In brief, the applied volume of Amberlite XAD-7 in the column was set to 150% of the theoretical binding capacity to ensure a sufficient removal of TnBP and Triton X-100. The resin column was equilibrated with 0.01 M phosphate buffer (Na_2_HPO_4_ and NaH_2_PO_4_) pH 7.1 containing 0.5 M NaCl until OD_280nm_ was < 0.02. Each S/D treated fecal virome (mixed by cage) was added separately to the column and the OD_280nm_ was measured to follow the concentration of proteins (expected viral particles and other metabolites > 30 kDa) and until OD_280nm_ was < 0.02. A 0.01 M phosphate buffer containing 1 M NaCl was used to release potential residual particles from the resin^101^. The removal of the solvent/detergent agents from the fecal virome yielded approx. 100 mL viral-flow-through from the column, which was concentrated to 0.5 mL using Centriprep Ultracel YM-30K units. The final product constituted the FVT-SDT treatment and was stored at -80 °C.

### Pyronin Y treated fecal virome (FVT-PyT)

Pyronin Y (Merck) is a strong, red-colored fluorescent compound. It has been reported to exhibit efficient binding to single-stranded and double-stranded RNA (ss/dsRNA), while its binding to single-stranded and double-stranded DNA (ss/dsDNA) is less effective^37,38^. The fecal filtrate was treated with 100µM pyronin Y and incubated at 40°C overnight to inactivate viral particles containing RNA genomes^30^. To remove the pyronin Y molecules that were not bound, the pyronin Y treated fecal filtrate suspensions were diluted in 50mL SM-buffer and subsequently concentrated to 0.5mL using Centriprep Ultracel YM-30K units. This process was repeated three times, resulting in a transparent appearance of the pyronin Y treated fecal filtrate, which constituted the FVT-PyT treatment and was stored at -80 °C.

### Chemostat propagated fecal virome (FVT-ChP)

The preparation of the chemostat propagated virome was performed as described in another study^31^. Briefly, anaerobic-handled mouse cecum content was utilized for chemostat propagation. The culture medium was formulated to resemble the low-fat (LF) diet (Research

Diets D12450J) provided to the donor mice as their feed, and growth conditions such as temperature (37°C) and pH (6.4) were set to simulate the environmental conditions present in the mouse cecum. The end cultures, which underwent fermentation with a slow dilution rate (0.05 volumes per hour), exhibited a microbial composition that resembled the initial microbial composition profile of the donor material^31^. These batches were combined to form the FVT-ChP treatment, and was stored at -80 °C.

### Fluorescence microscopy

Virus-like particle (VLP) counts were evaluated of all fecal viromes (unmodified FVT, FVT-SDT, FVT-ChP, and FVT-PyT, Fig. S1E) by epifluorescence microscopy using SYBR Gold staining (Thermo Scientific) as described online dx.doi.org/10.17504/protocols.io.bx6cpraw. The viral concentration was normalized using SM buffer to 2 x 10^9^ VLP/mL per treatment.

### Animal model design of diet-induced obesity

Forty-eight male C57BL/6NTac mice at 5 weeks old (Taconic Biosciences A/S, Lille Skensved, Denmark) were divided randomly (simple randomization) into 6 groups divided in 12 cages with 4 mice per cage: lean control, obese control, unmodified FVT (as the control for modified FVTs), FVT-ChP, FVT-SDT, and FVT-PyT (Fig. 6). Each group was represented by 8 mice where the single animals were interpreted as the experimental unit. Mice were housed with 2 cages per group to account for potential cage effect bias. Decision on sample size (n = 8) was based on a previous publication where we validated the suitable sample size in measuring prediabetic symptoms by either HbA1c levels or oral glucose tolerance test (OGTT)^95^, which was further supported by a subsequent study^25^. Only male C57BL/6NTac mice were included since female mice are highly protected against diet-induced obesity^42^. For 18 weeks, mice were fed *ad libitum* high-fat diet (Research Diets D12492), except lean controls that were fed a low-fat diet (Research Diets D12450J). After 6 weeks on their respective diets, mice were administered 0.15 mL of the unmodified FVT, FVT-ChP, FVT-SDT, or FVT-PyT by oral gavage twice with one week of interval (study weeks 7 and 8). Obese and lean controls received 0.15 mL SM buffer as sham. The titre of the applied FVT virome was approximately 2 × 10^9^ VLP/mL (Fig. S10A-D). The mice were subjected to an OGTT at week 13 and 18 of the study (18 and 23 weeks old) by blinded personnel, and food intake and mouse weight were monitored frequently to evaluate sudden diabetes associated weight-loss and behavioral changes (only the ear-tagged mouse ID was available). Author TSR was aware at all stages of the group allocation of the cages, but not the ear-tagged mouse ID, while the animal caretakers were blinded at all stages, thus TSR handed the FVT treatments or saline to the animal caretakers. OGTT outliers were removed if 2 of 3 parameters were applicable: 1) tAUC > 2000 mg/dL, fasting glucose > 11.5 mM, or 3) loss of 5-10% in body weight. Treatments, handling, and sampling of cages (cage 1-12) were performed in the order as lean control, obese control, unmodified FVT, FVT-ChP, FVT-SDT, and FVT-PyT (cage 1-6) and repeated with cage 7-12 in same group order. The location of the cages had similar expose to light, noise, distance to ceiling, floor, and entrance. Blood serum, epididymal white adipose tissue (eWAT), the mesenteric lymph node (MLN), liver tissue, intestinal content from the cecum and colon, and tissue from the colon and ileum was sampled at termination at study week 18 (23 weeks old) and stored at -80°C until downstream analysis. Liver tissue was fixated in 10% neutral-buffered formalin (Sarstedt Formalin System) for histological analysis and stored at room temperature until further processed. All procedures regarding the handling of animals included in the diet-induced obesity model were carried out in accordance with the Directive 2010/63/EU and the Danish Animal Experimentation Act with the license ID: 2017-15-0201-01262 C1, and the and housing and enrichment conditions were as earlier described^48^.

**Fig. 6:**
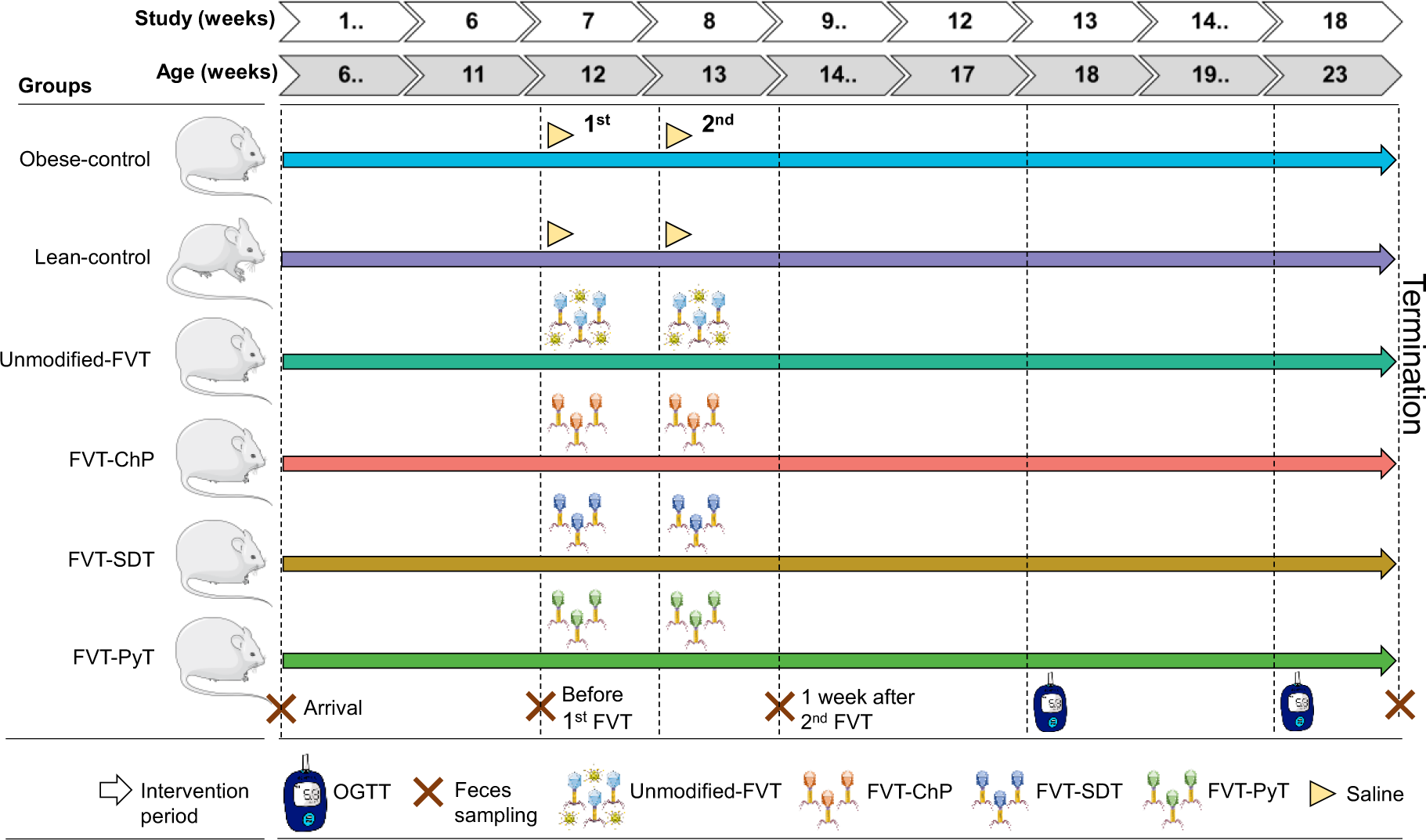
Overview of the animal study design. Forty-eight male C57BL/6NTac mice (5 weeks old) were divided into six groups. The mice were fed with either a high-fat or low-fat diet. The mice were treated with different FVTs or saline, after 6 and 7 weeks on their respective diets. The FVT treatments represented an unmodified FVT (containing all viruses but limited number of viable bacteria), chemostat propagated fecal virome nearly depleted from eukaryotic viruses by dilution (FVT-ChP), solvent-detergent treated FVT to inactivate enveloped viruses (FVT-SDT), and pyronin Y treated FVT to inactive RNA viruses, and a saline solution for the obese and lean control. Oral glucose tolerance tests (OGTT) were measured twice (18 and 23 weeks old). The crosses mark the timepoint of feces samples that were used for gut microbiome analysis.

### Cytokine analysis

Serum samples were diluted 1:2 and analyzed for IFN-γ, GM-CSF, IL-15, IL-6, IL-10, KC/GRO, MIP-2 TNF-α, IL-17A/F, and IL-22 in a customized metabolic group 1 U-PLEX (Meso Scale Discovery, Rockville, ML) according to manufacturer’s instructions. Samples were read using the MESO QuickPlex SQ 120 instrument (Meso Scale Discovery) and concentrations were extrapolated from a standard curve using their software Discovery Workbench v.4.0 (Meso Scale Discovery). Measurements out of detection range were assigned the value of lower or upper detection limit. The cytokine analysis was performed by a blinded investigator.

### Histology

Formalin-fixed liver biopsies were embedded in paraffin, sectioned, and stained with hematoxylin & eosin for histopathological evaluation. We used a cumulative semi-quantitative MAFLD activity score (MAS) comprising the following three histological features: Steatosis (0-3), immune cell margination and infiltration (0-2), and hepatocellular ballooning (0-2) with higher values corresponding to increased dissemination. The histological assessment was performed by a blinded investigator.

### Cell isolation and fluorescence-activated cell sorting (FACS)

Directly after sacrificing the mice, the MLN and eWAT were placed in ice cold PBS. Single-cell suspensions were prepared by disrupting the lymph node between two microscope glasses and passing it through a 70 μm nylon mesh. eWAT cells were isolated as previously described^102^. After washing in PBS and resuspension, 1 x 10^6^ cells were surface stained for 30 min with antibodies for PerCP-Cy5.5 conjugated CD11c, PE-conjugated CD19, APC-conjugated F4.80, and FITC-conjugated CD45 (all antibodies were purchased from eBiosciences) for the detection of the different antigen-presenting cells. For the detection of T cell subsets, 1x10^6^ cells were initially surface stained for 30 min with FITC-conjugated CD8α, PE-conjugated CD4, and APC-conjugated TCRαβ (ebiosciences). Specifically for the differentiation of memory vs. naïve T cells, the cells were stained with FITC-conjugated CD62l, PE-conjugated CD44, PerCP-Cy5.5-conjugated CD4, and APC-conjugated CD8α (ebiosciences). Analysis was performed using an Accuri C6 flow cytometer (Accuri Cytometers, now BD Biosciences) and accompanying software BD Accuri C6 software version 264.21. The FACS measures were performed by blinded investigators.

### Pre-processing of fecal samples for separation of viruses and bacteria

The gut bacterial and viral composition of the mice were investigated at respectively four and three time points: upon arrival at our housing facilities (5 weeks old), before 1^st^ FVT (11 weeks old, but only the bacteriome), one week after the 2^nd^ FVT (13 weeks old), and at termination (23 weeks old), which represented in total 192 fecal samples. Separation of viruses and bacteria from fecal samples generated a fecal pellet and fecal supernatant by centrifugation and 0.45 µm filtering as described previously^48^, however, the volume of fecal homogenate was adjusted to 5 mL SM buffer.

### Bacterial DNA extraction, sequencing, and pre-processing of raw data

The DNeasy PowerSoil Pro Kit (Qiagen) was used to extract bacterial DNA from the fecal pellet by following the instructions of the manufacturer and was performed by author XM who at this stage was blinded. The final purified DNA was stored at -80°C, and the DNA concentration was determined using Qubit HS Assay Kit (Invitrogen) on the Qubit 4 Fluorometric Quantification device (Invitrogen). The bacterial community composition was determined by Illumina NextSeq-based high-throughput sequencing (HTS) of the 16S rRNA gene V3-region, as previously described^48^. Quality control of reads, de-replicating, purging from chimeric reads and constructing zOTU was conducted with the UNOISE pipeline and taxonomically assigned with Sintax. Taxonomical assignments were obtained using the EZtaxon for 16S rRNA gene database. Code describing this pipeline can be accessed in github.com/jcame/Fastq_2_zOTUtable. The average sequencing depth after quality control (Accession: PRJEB58786, available at ENA) for the fecal 16S rRNA gene amplicons was 67,454 reads (min. 12,790 reads and max. 295,746 reads).

### Viral RNA/DNA extraction, sequencing and pre-processing of raw data

The sterile filtered fecal supernatant was concentrated using centrifugal filters Centrisart with a filter cut-off at 100 kDA (Sartorius) by centrifugation centrifuged at 1,500 x g at 4°C (dx.doi.org/10.17504/protocols.io.b2qaqdse). Fecal supernatant (140 µL) was treated with 5 units of Pierce™ Universal Nuclease (ThermoFisher Scientific) for 10 minutes at room temperature prior to viral DNA extraction to remove free DNA/RNA molecules. The viral DNA/RNA was extracted from the fecal supernatants with Viral RNA mini kit (Qiagen) as previously described^48^, and was performed by author XM who at this stage was blinded. Reverse transcription was performed with SuperScript VILO Master mix by following the instructions of the manufacturer and subsequently cleaned with DNeasy blood and tissue kit (Qiagen) by only following step 3-8 in the manual. In brief, the DNA/cDNA samples were mixed with ethanol, bound to the silica filter, washed two times, and eluted with 40 µL elution buffer. Multiple displacement amplification (MDA, to include ssDNA viruses) using GenomiPhi V3 DNA amplification kit (Cytiva) and sequencing library preparation using Nextera XT kit was performed as previously described^48^, and sent for sequencing using the NovaSeq platform (NovoGene). The average sequencing depth of raw reads (Accession: PRJEB58786, available at ENA) for the fecal viral metagenome was 13,145,283 reads (min. 693,882 reads and max. 142,821,858 reads. Using Trimmomatic v0.35, raw reads were trimmed for adaptors and low quality sequences (<95 % quality, <50nt) were removed. High-quality reads were de-replicated and checked for the presence of PhiX control using BBMap (bbduk.sh) (https://www.osti.gov/servlets/purl/1241166). Virus-like particle-derived DNA sequences were subjected to within-sample de-novo assembly-only using Spades v3.13.1 and the contigs with a minimum length of 2,200 nt, were retained. Contigs from all samples were pooled and dereplicated by chimera-free species-level clustering at ∼95 % identity using the script described in^103^, and available at https://github.com/shiraz-shah/VFCs. Contigs were classified as viral by VirSorter2^104^ ("full" categories | dsDNAphage, ssDNA, RNA, Lavidaviridae, NCLDV | viral quality = 1), VIBRANT^105^ (High-quality | Medium-quality | Complete), CheckV^106^ (High-quality | Medium-quality | Complete), and VirBot^107^. Any contigs not classified as viral by any of the four software’s were discarded. The taxonomical categories of "Other/Remaining taxa", "Unclassified virus", and "Unknown" that are used in the different figures are different entities. "Other/Remaining taxa" encompasses all remaining low abundance taxa not depicted in the plot. "Unknown" refers to contigs that may be viruses but lack specific data records confirming their viral origin, and "Unclassified virus" represents viruses that have been identified as having viral origin but could not be further classified. Taxonomy was inferred by blasting viral ORFs against a database of viral proteins created from the following: VOGDB v217 (vogdb.org), NCBI (downloaded 14/10/2023), COPSAC^103^and an RNA phage database^108^, selecting the best hits with a minimum e-value of 10e^-6^. Phage-host predictions were done with IPhoP^109^, which utilizes a combination of other host predictors. Following assembly, quality control, and annotations, reads from all samples were mapped against the viral (high-quality) contigs (vOTUs) using bowtie2^110^ and a contingency table of contig-length and sequencing-depth normalized reads, here defined as vOTU-table (viral contigs). Code describing this pipeline can be accessed in https://github.com/frejlarsen/vapline3. Mock phage communities (phage C2, T4, phiX174, MS2, and Phi6, Table S3) were used as positive controls (normalized to ∼10^6^ PFU/mL for each phage) for virome sequencing to validate the sequencing protocol’s ability to include the different genome types of ssDNA, dsDNA, ssRNA, and dsRNA.

### Bioinformatic analysis and statistics

Initially, the dataset was purged for zOTU’s/viral contigs, which were detected in less than 5% of the samples, but the resulting dataset still maintained 99.8% of the total reads. R version 4.3.0 was used for subsequent analysis and presentation of data. A minimum threshold of sequencing reads for the bacteriome and virome analysis was set to 2,000 reads and 15,000 reads, respectively. The main packages used were phyloseq^111^, vegan^112^, DESeq2^113^, ampvis2^114^(not yet peer-reviewed), ggpubr, psych, igraph, ggraph, pheatmap, ComplexHeatmap, and ggplot2. The contamination of viral contig was removed by read count detected in negative controls through R package microDecon^115^ (runs = 1, regressions = 1), and 41.5% of entries were removed. Cumulative sum scaling (CSS) normalization was performed using the R software using the metagenomeSeq package. α-diversity analysis was based on raw read counts and statistics were based on ANOVA. β-diversity was represented by Bray-Curtis dissimilarity and statistics were based on pairwise PERMANOVA corrected with FDR (false discovery rate). DESeq2 was used to identify differential microorganisms on the summarized bacterial species level and viral contigs (vOTUs) level. The correlation network of bacterial association with phenotypic and immunologic variables, and the correlation heatmap between bacterial zOTUs and viral contigs (vOTUs) were calculated using pairwise Spearman’s correlations and corrected with FDR. The non-parametric two-side Wilcoxon rank-sum tests were adopted for analysis of the phenotypic variables, cytokine levels, immune cell levels, Shannon diversity index (α-diversity), PERMANOVA of β-diversity, and abundance of single bacterial genus, the comparison was conducted between FVT treatments and obese control mice, with FDR correction adopted, the comparison between obese and lean control mice was also conducted.

## Acknowledgments

We thank the animal caretakers Helene Farlov and Mette Nelander at Section of Experimental Animal Models (University of Copenhagen, Denmark) for taking care of the animals during the study and assisting with the animal handling. A thanks to PhD fellow Kaare Dyekær Tranæs (Copenhagen Prospective Studies on Asthma in Childhood, Copenhagen University Hospital) for contributing with internal review and proof-reading of the manuscript.

## Author contributions

TSR and DSN conceived the research idea and designed the study. TSR, SBL, LSFZ, CHFH, and AKH conducted, monitored, and supervised the animal experiments. TSR, SBL, DSN, SA, and KA prepared and designed the methodology of the chemostat-propagated virome (FVT-ChP). TSR and SBL prepared and designed the methodology of the unmodified FVT, solvent-detergent (FVT-SDT) and pyronin Y treated FVT (FVT-PyT) and evaluated viral-like-particle count with fluorescence microscopy. XM and TSR performed nucleotide extractions and library preparation for sequencing. LSFZ and CHFH conducted immune cell count and cytokine profile analysis. AB prepared and scored the liver histology. TSR, SBL, XM, FL, JLCM, and DSN contributed to the bioinformatic analysis and interpretation. TSR, SBL, XM, CHFH, LSFZ, DSN, AKH, and AB analyzed and interpreted the phenotypical measures of the mice. TSR and DSN supervised the study. DSN was responsible for the funding. XM and TSR wrote the first draft of the manuscript. All authors critically revised and approved the final version of the manuscript.

## Ethics declarations

### Animal welfare

All procedures involving handling of animals included in the diet-induced obesity model (license ID: 2017-15-0201-01262 C1) and donor animals (license ID: 2012-15-2934-00256) were approved by the Animal Experiments Inspectorate (Ministry of Food, Agriculture, and Fisheries of Denmark) and conducted in accordance with Directive 2010/63/EU and the Danish Animal Experimentation Act.

### Inclusion

All collaborators in this study have met the authorship criteria mandated by Nature Portfolio journals and have been included as authors because their participation was crucial for designing and conducting the study. Roles and responsibilities were agreed upon among collaborators before the research commenced. This work encompasses findings that are locally and internationally relevant. The research was not subject to severe restrictions or prohibitions in the researchers’ setting.

## Data availability statement

All data associated with this study are present in the paper or in the Supplementary Materials, or at https://github.com/MaoAria15/DIO. All 529 sequencing datasets are available in the ENA database under accession number PRJEB58786.

## Code availability statement

The codes used for our analyses are available at https://github.com/MaoAria15/DIO, https://github.com/frejlarsen/vapline3, and https://github.com/jcame/Fastq_2_zOTUtable.

## Supplementary materials

Supplementary materials Peer Review file Reporting Summary ARRIVE Essential 10 SAGER Checklist

