## Supplemental materials for "Transfer of modified fecal viromes improve blood glucose regulation and alleviates symptoms of metabolic dysfunction-associated fatty liver disease in an obesity male mouse model"

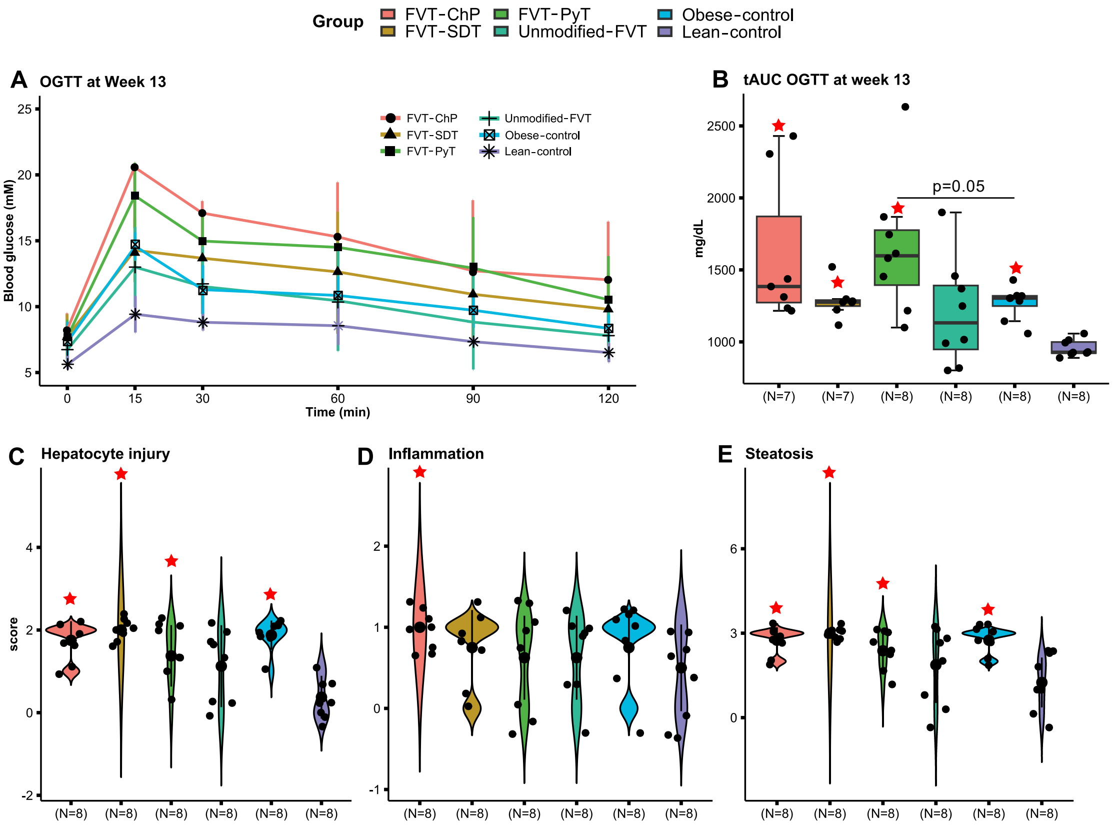


Fig. S1: Overview of mouse phenotypic characteristics after being treated with the different FVTs. A) Total area under the curve (tAUC) and B) line plot of OGTT measured over 120 min of the different FVT treatments at study week 13 (18 weeks old), and C) to E) sub-categories of MAFLD activity score evaluating hepatocyte injury, inflammation, and steatosis at study week 18 (23 weeks old). The labels of *, **, *** between the FVT treatments and obese control mice represent adjusted p < 0.05, 0.01, 0.001 (two-side Wilcoxon rank-sum test with FDR correction); the unadjusted p is presented in the figure when it is less than 0.05. The ★ label on top of the boxplot of the treatment represents a p < 0.05 between the treatment and the lean control mice (two-side Wilcoxon rank-sum test). Combined boxplots distributions include median, min, max, 25 and 75 percentiles, and outliers (more than 1.5 IQR).

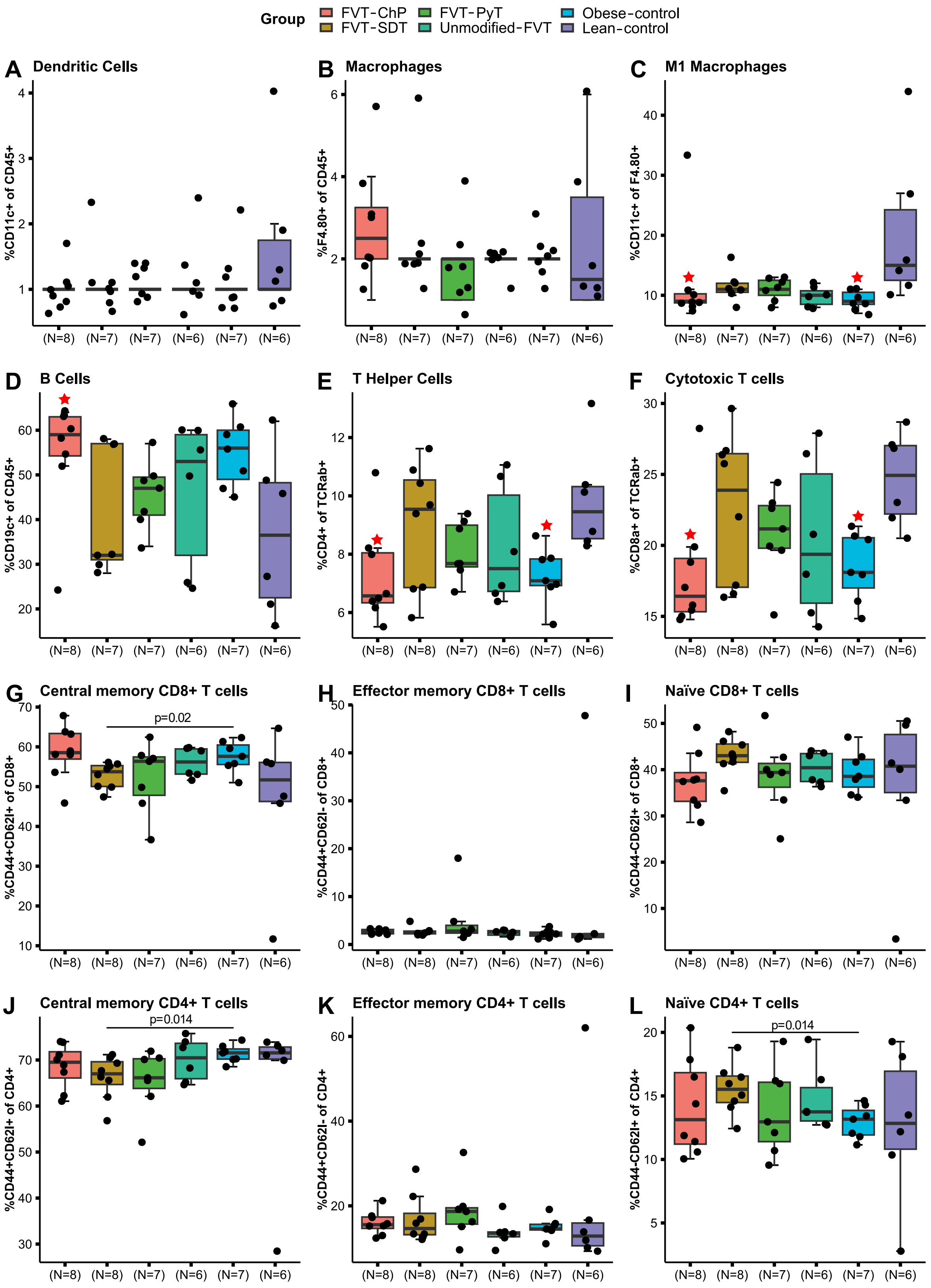
Group
FVT-ChP
FVT-SDT
FVT-PyT
Unmodified-FVT
Obese-control
Lean-control
A Dendritic Cells
%CD11c+ of CD45+
B Macrophages
%F4.80+ of CD45+
C M1 Macrophages
%CD11c+ of F4.80+
D B Cells
%CD19c+ of CD45+
E T Helper Cells
%CD4+ of TCRab+
F Cytotoxic T cells
%CD8a+ of TCRab+
G Central memory CD8+ T cells
p=0.02
%CD44+CD62l+ of CD8+
H Effector memory CD8+ T cells
%CD44+CD62l- of CD8+
I Naïve CD8+ T cells
%CD44-CD62l+ of CD8+
J Central memory CD4+ T cells
p=0.014
%CD44+CD62l+ of CD4+
K Effector memory CD4+ T cells
%CD44+CD62l- of CD4+
L Naïve CD4+ T cells
p=0.014
%CD44-CD62l+ of CD4+

10

Fig. S2: Proportions of immune cells in the mesenteric lymph node (MLN) at study termination (23 weeks old). A) to L) show the overall FACS profile in the MLN tissue of the different treatments. The labels of *, **, *** between the FVT treatments and obese control mice represent adjusted $p < 0.05, 0.01, 0.001$ (two-side Wilcoxon rank-sum test with FDR correction), the unadjusted p is presented in figure when it is less than 0.05. The ★ label on top of the boxplot of the treatment represents a $p < 0.05$ between the treatment and the lean control mice (two-side Wilcoxon rank-sum test). Combined boxplots distributions include median, min, max, 25 and 75 percentiles, and outliers (more than 1.5 IQR).

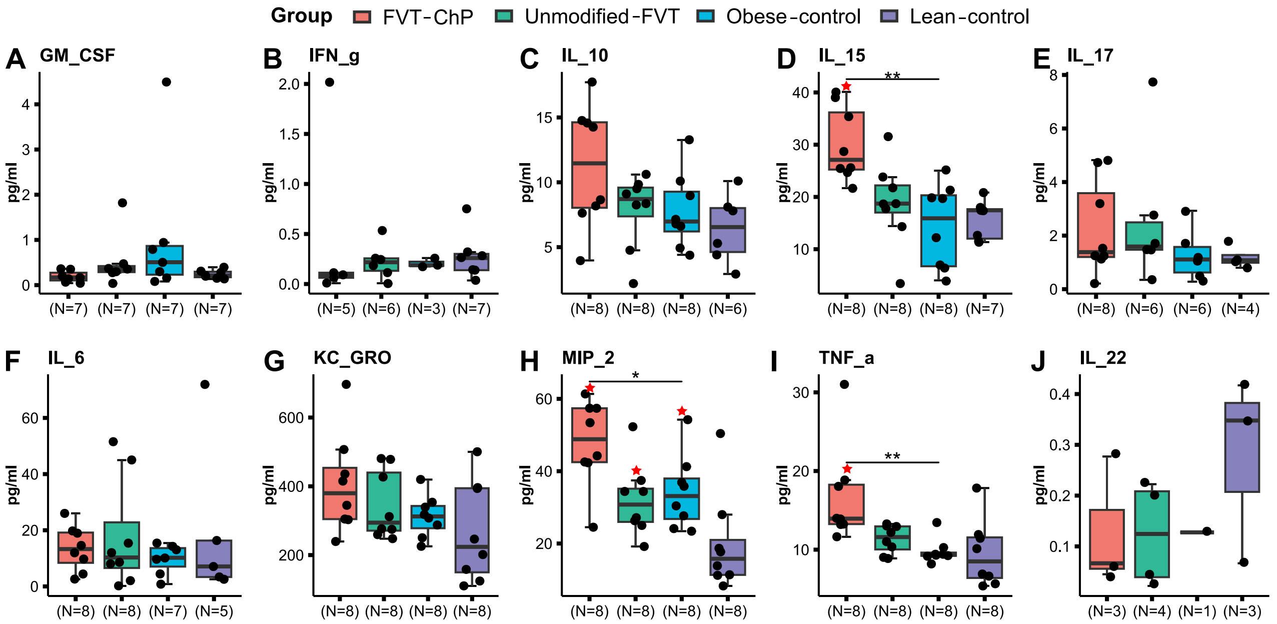


Fig. S3: Cytokine profiles of blood serum sampled at termination (23 weeks old). A) to J) are showing the overall cytokine profile in the mouse blood serum of mice treated with FVT-ChP, unmodified FVT, and the two dietary control groups. The labels of *, **, *** between the FVT treatments and obese control mice represent adjusted $p < 0.05, 0.01, 0.001$ (two-side Wilcoxon rank-sum test with FDR correction), the unadjusted p is presented in figure when it is less than 0.05. The ★ label on top of the boxplot of the treatment represents a $p < 0.05$ between the treatment and the lean control mice (two-side Wilcoxon rank-sum test). Combined boxplots distributions include median, min, max, 25 and 75 percentiles, and outliers (more than 1.5 IQR).

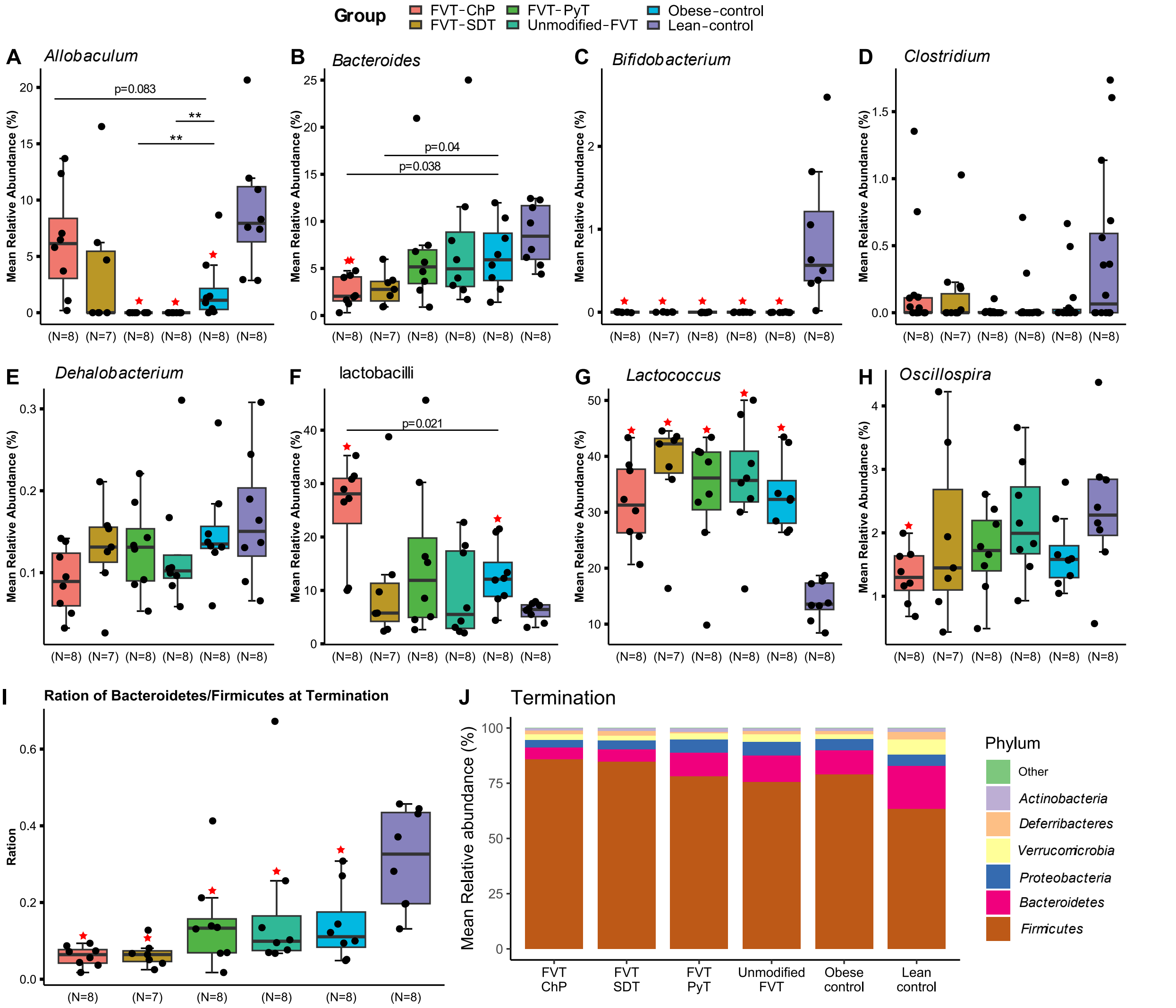


Fig. S4: Relative bacterial abundance at termination (23 weeks old). A) – H) Boxplots showing the relative abundance of selected bacterial taxa. I) Boxplots of the ratio between *Bacteroidetes*:*Firmicutes* that were observed with the different treatment groups. J) Bar plot of phyla distribution based on treatment group. The labels of *, **, *** between the FVT treatments and obese control mice represent adjusted p < 0.05, 0.01, 0.001 (two-side Wilcoxon rank-sum test with FDR correction), the unadjusted p is presented in figure when it is less than 0.05. The ★ label on top of the boxplot of the treatment represents a p < 0.05 between the treatment and the lean control mice (two-side Wilcoxon rank-sum test). Combined boxplots distributions include median, min, max, 25 and 75 percentiles, and outliers (more than 1.5 IQR).

See the additional high-resolution PDF Fig. S5

Fig. S5: Heatmap highlighting significant ($p < 0.05$, FDR correction) differences in the differential abundance of viral contigs at termination (23 weeks old).

See the additional high-resolution PDF Fig. S6

Fig. S6: Pairwise Spearman’s correlation between relative abundance of bacterial zOTUs (relative abundance > 0.1%) and viral vOTUs (relative abundance > 0.1%), with the FDR correction for multi-comparison (study week 18, 23 weeks old).

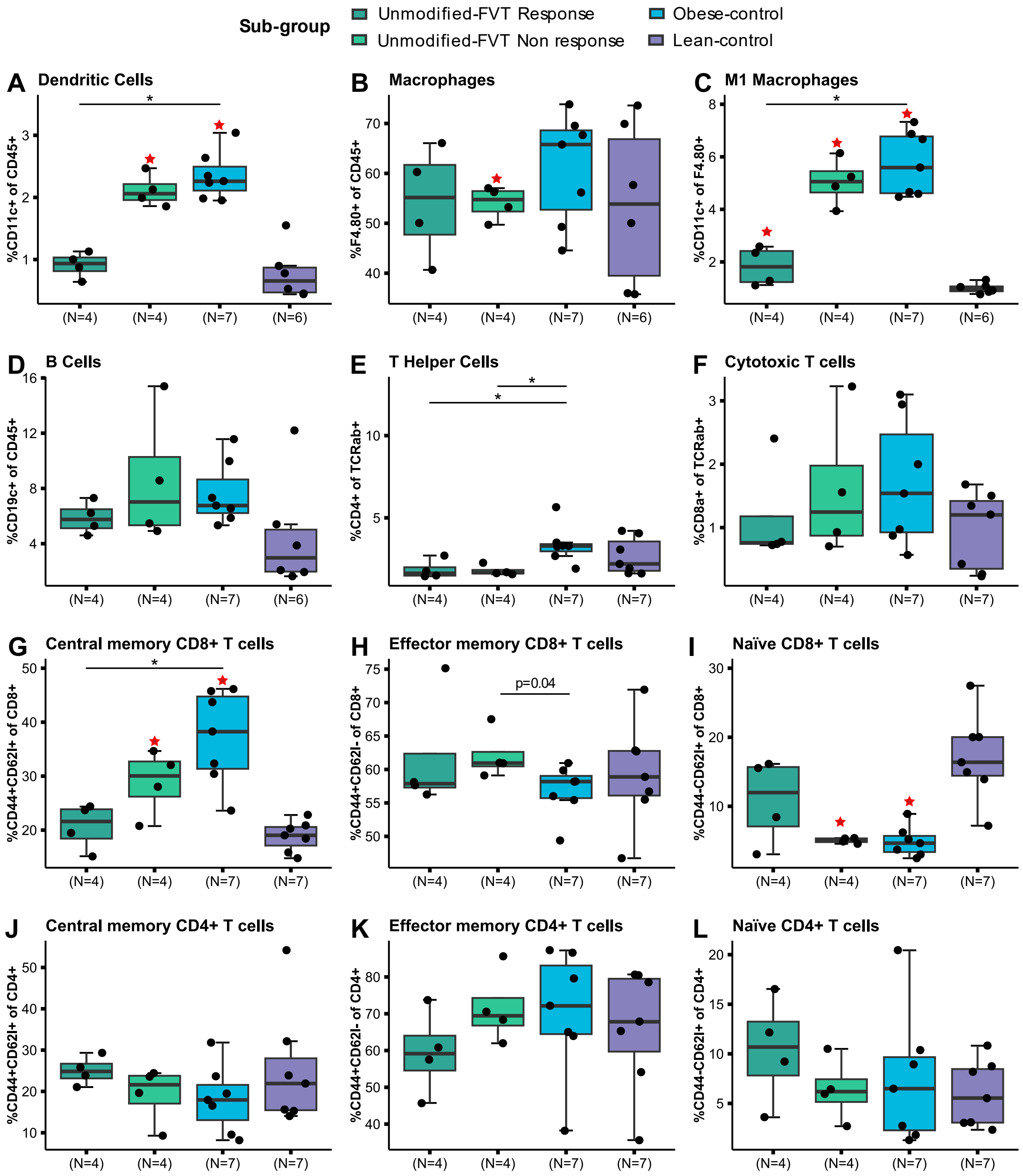


Fig. S7: Proportions of immune cells in adipose tissue of the unmodified FVT treatment sub-test at study termination (23 weeks old). A) to L) are showing the overall fluorescence-activated cell sorter (FACS) profile in the mouse adipose tissue. The labels of *, **, *** between the FVT treatments and obese control mice represent adjusted $p < 0.05, 0.01, 0.001$ (two-side Wilcoxon rank-sum test with FDR correction), the unadjusted p is presented in figure when it is less than 0.05. The ★ label on top of the boxplot of the treatment represents a $p < 0.05$ between the treatment and the lean control mice (two-side Wilcoxon rank-sum test). Combined boxplots distributions include median, min, max, 25 and 75 percentiles, and outliers (more than 1.5 IQR).

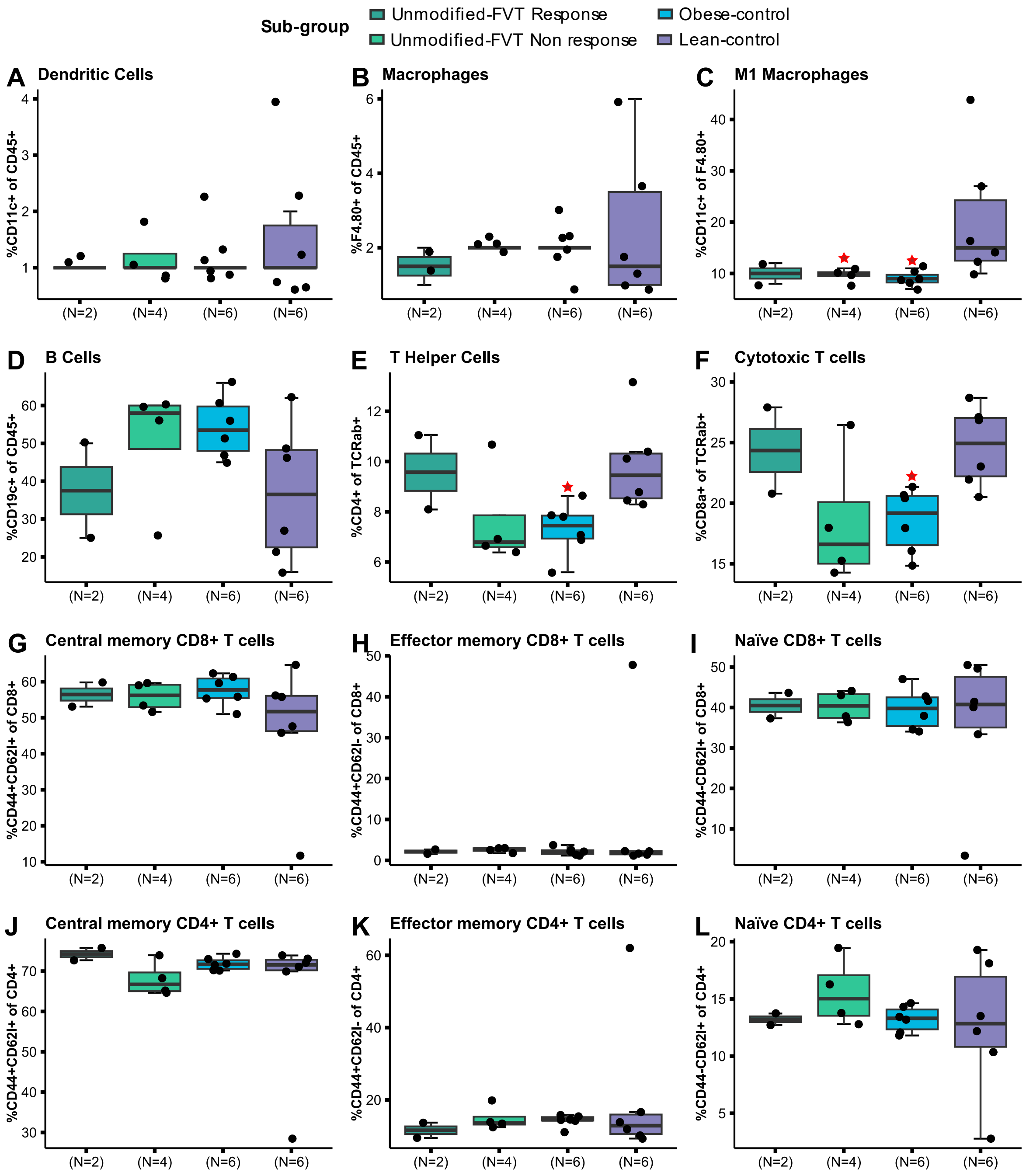


Fig. S8: Proportions of immune cells in mesenteric lymph node (MLN) of the unmodified FVT treatment sub-test at study termination (23 weeks old). A) to L) show the overall FACS profile in the MLN tissue of the different treatments. The labels of *, **, *** between the FVT treatments and obese control mice represent adjusted $p < 0.05$, 0.01, 0.001 (two-side Wilcoxon rank-sum test with FDR correction), the unadjusted p is presented in figure when it is less than 0.05. The ★ label on top of the boxplot of the treatment represents a $p < 0.05$ between the treatment and the lean control mice (two-side Wilcoxon rank-sum test). Combined boxplots distributions include median, min, max, 25 and 75 percentiles, and outliers (more than 1.5 IQR).

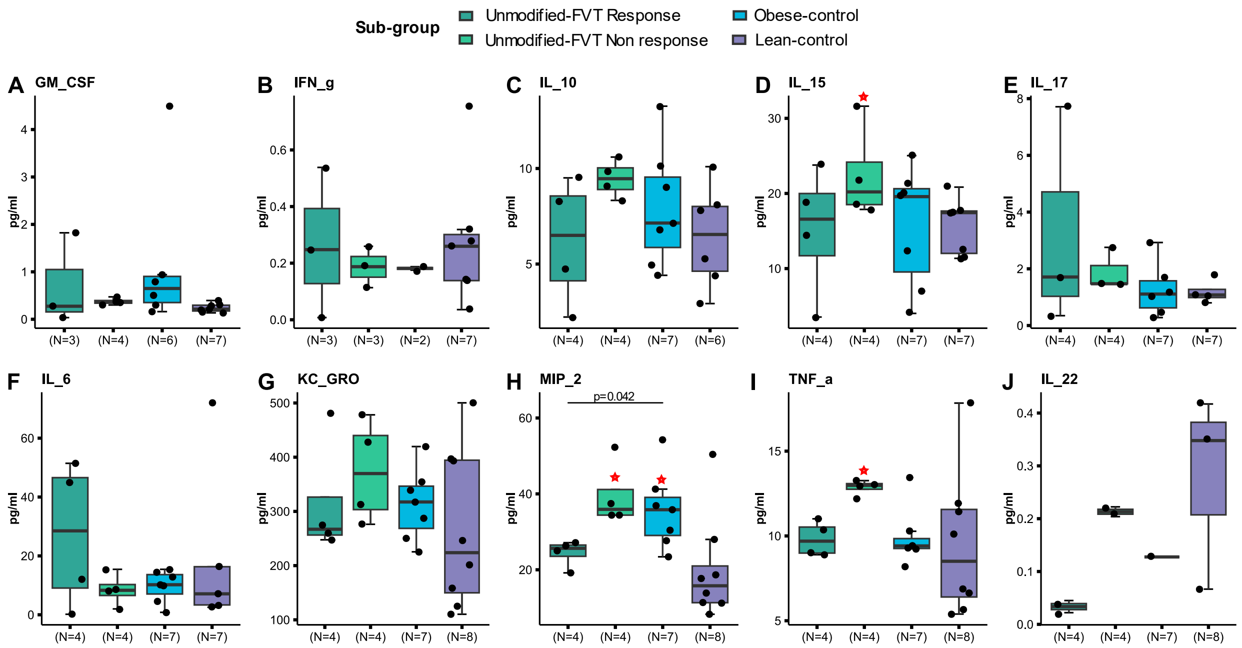


Fig. S9: Cytokine profiles of blood serum of the unmodified FVT treatment sub-test at study termination (23 weeks old). A) to J) are showing the overall cytokine profile in the mouse blood serum of mice that responded or not responded on the unmodified FVT, and the two dietary control groups. The labels of *, **, *** between the FVT treatments and obese control mice represent adjusted $p < 0.05, 0.01, 0.001$ (two-side Wilcoxon rank-sum test with FDR correction), the unadjusted p is presented in figure when it is less than 0.05. The ★ label on top of the boxplot of the treatment represents a $p < 0.05$ between the treatment and the lean control mice (two-side Wilcoxon rank-sum test). Combined boxplots distributions include median, min, max, 25 and 75 percentiles, and outliers (more than 1.5 IQR).

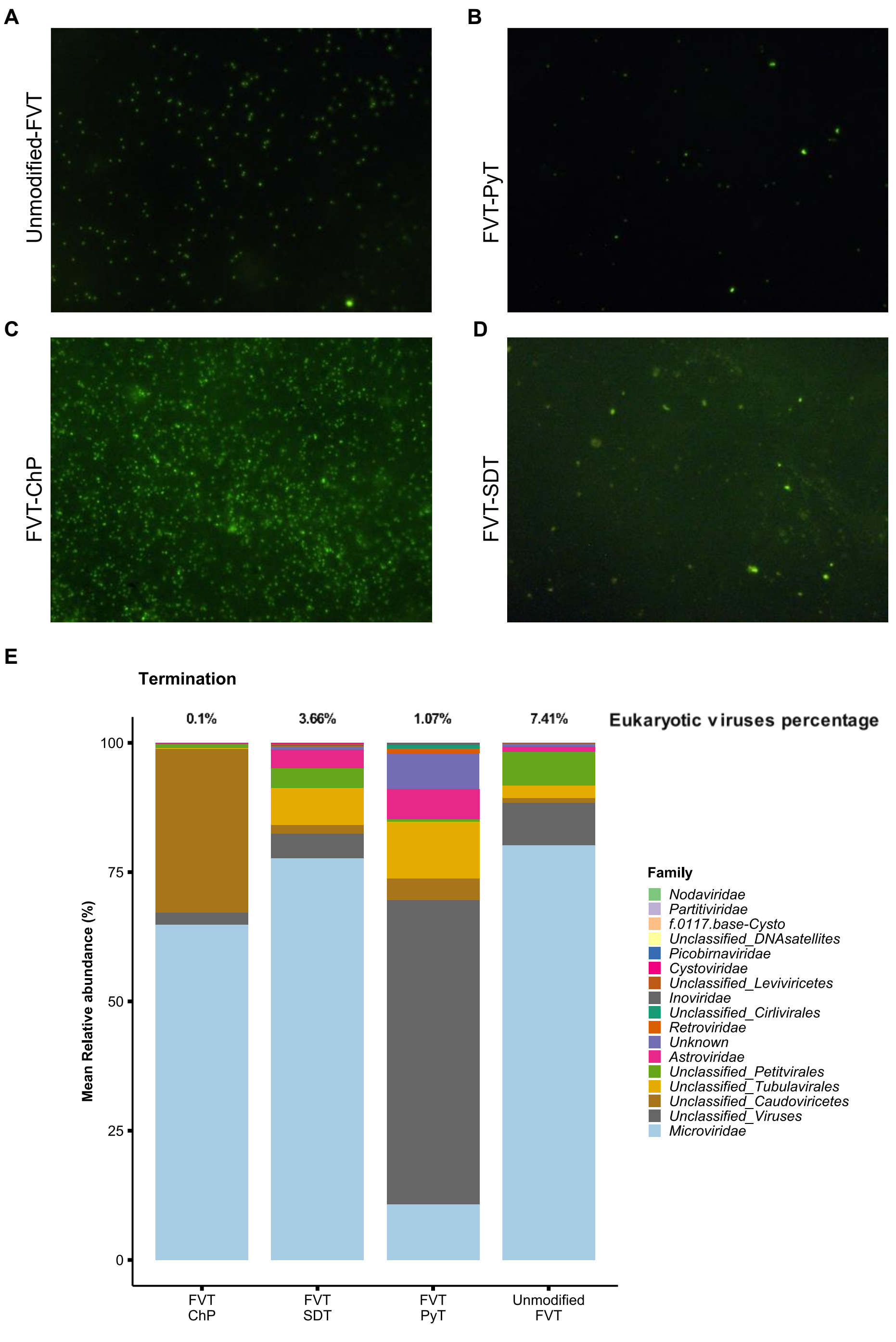


Fig. S10: Overview of the applied FVT viromes. A) – D) Epifluorescence microscopy images of the different applied FVT viromes (before being diluted to similar VLP/mL concentrations) that were stained with SYBR Gold to count VLP/mL. E) Viral taxonomy composition bar plot of the four applied FVTs on the family level including the relative abundance in percentages of eukaryotic viruses in each FVT treatment.

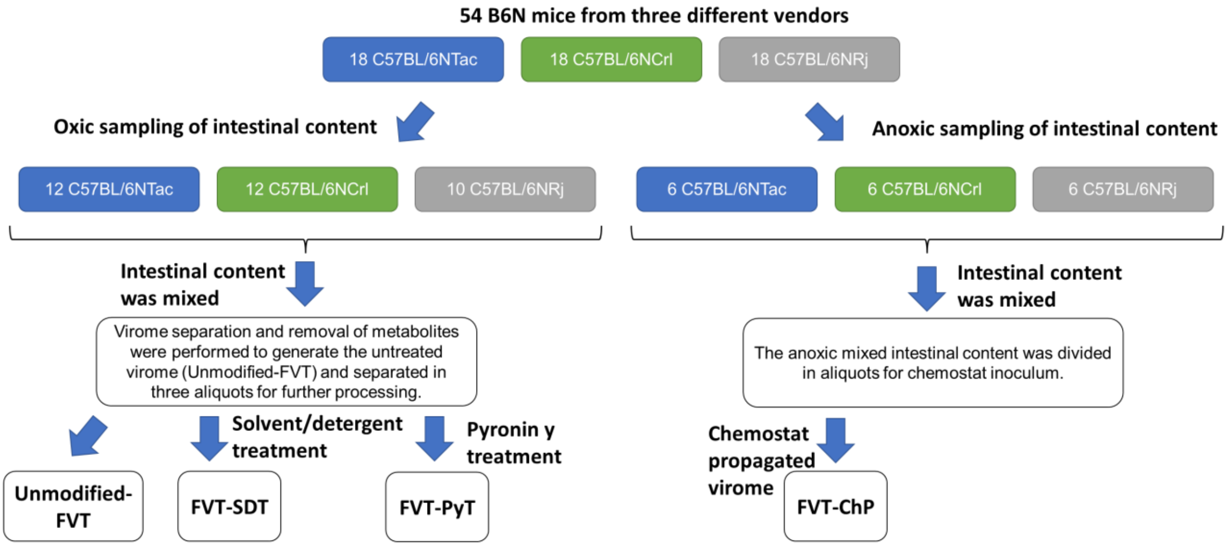


Fig. S11: Flow diagram illustrating the origin of the intestinal donor content and the processing steps to generate the FVTs with different methodologies. Initially, 54 mice from 3 different vendors were sacrificed and their intestinal content from cecum and colon was collected, however, 34 mice were sampled in oxic conditions, while 18 mice were sampled in anoxic conditions to maintain the viability of the strict anaerobic bacterial GM members. Two C57BL/6NRj mice were euthanized due to malocclusion-associated malnutrition. The atmospheric conditions were maintained throughout the process. Regardless of the vendor, the intestinal content was mixed. Virome separation was performed for the oxic-handled fecal mixture and was divided into three aliquots that represented the untreated fecal virome (unmodified FVT), further processing with solvent/detergent treatment (FVT-SDT) or pyronin Y treatment (FVT-PyT). Anoxic handled fecal mixture was used as inoculum for virome propagation in a chemostat setup (FVT-ChP). Also, the FVT-ChP underwent virome separation to remove most metabolites as well as bacteria and other large microbes.

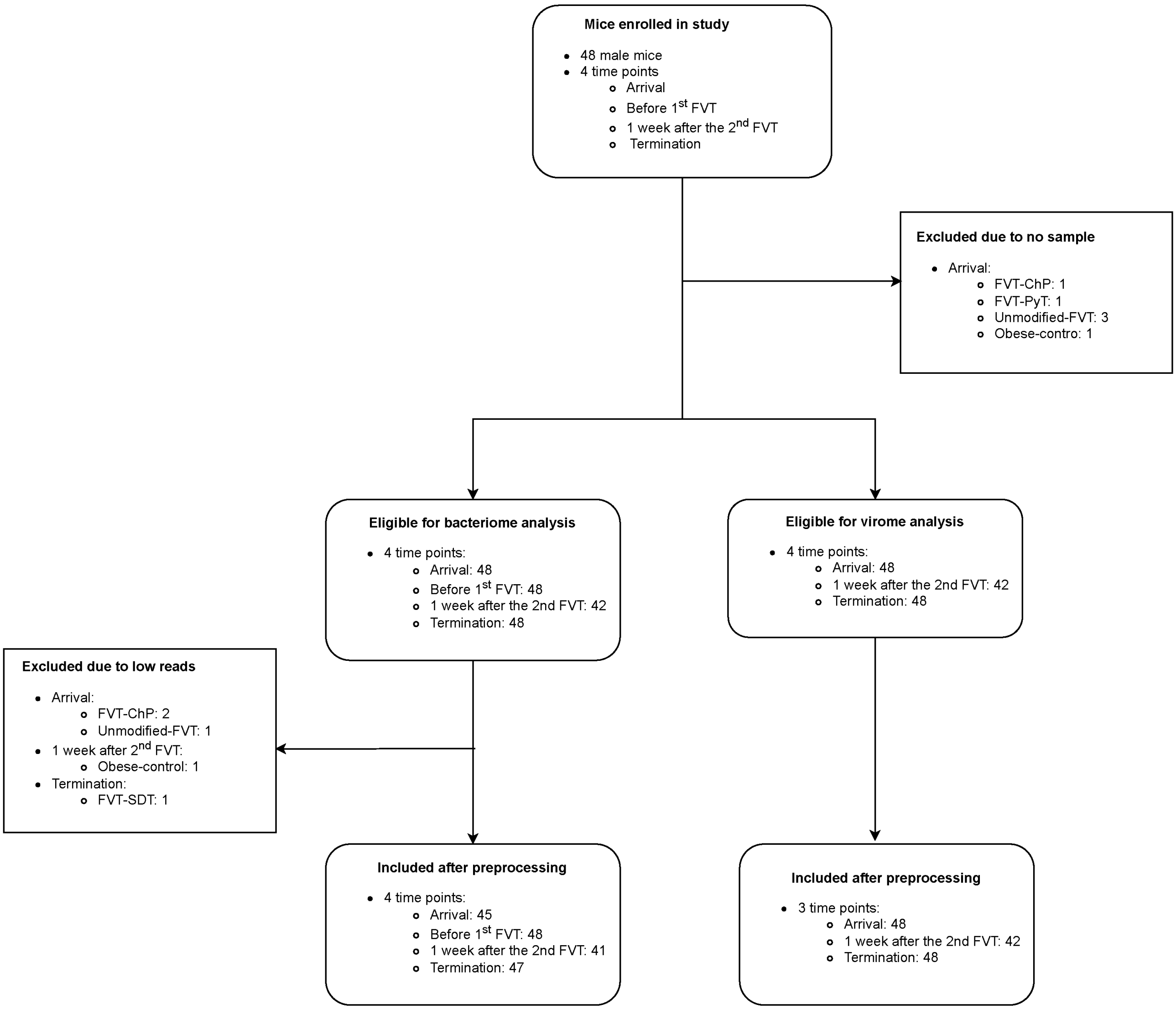


Fig. S12: Flowchart of fecal samples of mice inclusion and exclusion for bacteriome and virome analysis.

Table S1: Overview of the oral glucose tolerance (OGTT) statistics at each time point at study week 13 (18 weeks old). Comparison was done with Wilcoxon rank-sum test (two-side) with FDR correction which was only conducted between the FVT treatments and obese control mice.

| | **Group1** | **Group2** | **n1** | **n2** | **p** | **p.adj** |
|---|---|---|---|---|---|---|
| **t0** | FVT-ChP | Obese-control | 8 | 8 | 0.207 | 0.528 |
| | FVT-SDT | Obese-control | 8 | 8 | 0.399 | 0.528 |
| | FVT-PyT | Obese-control | 8 | 8 | 0.528 | 0.528 |
| | Unmodified-FVT | Obese-control | 8 | 8 | 0.429 | 0.528 |
| | Obese-control | Lean-control | 8 | 8 | 0.01 | - |
| | Lean-control | FVT-ChP | 8 | 8 | 0.006 | - |
| | Lean-control | FVT-SDT | 8 | 8 | 0.002 | - |
| | Lean-control | FVT-PyT | 8 | 8 | 0.012 | - |
| | Lean-control | Unmodified-FVT | 8 | 8 | 0.093 | - |
| **t15** | FVT-ChP | Obese-control | 8 | 8 | 0.002 | 0.006 |
| | FVT-SDT | Obese-control | 8 | 8 | 0.4 | 0.4 |
| | FVT-PyT | Obese-control | 8 | 8 | 0.05 | 0.1 |
| | Unmodified-FVT | Obese-control | 8 | 8 | 0.279 | 0.372 |
| | Obese-control | Lean-control | 8 | 8 | 0.00016 | - |
| | Lean-control | FVT-ChP | 8 | 8 | 0.012 | - |
| | Lean-control | FVT-SDT | 8 | 8 | 0.000311 | - |
| | Lean-control | FVT-PyT | 8 | 8 | 0.003 | - |
| | Lean-control | Unmodified-FVT | 8 | 8 | 0.015 | - |
| **t30** | FVT-ChP | Obese-control | 8 | 8 | 0.052 | 0.185 |
| | FVT-SDT | Obese-control | 8 | 8 | 0.442 | 0.589 |
| | FVT-PyT | Obese-control | 8 | 8 | 0.093 | 0.185 |
| | Unmodified-FVT | Obese-control | 8 | 8 | 0.875 | 0.875 |
| | Obese-control | Lean control | 8 | 8 | 0.003 | - |
| | Lean-control | FVT-ChP | 8 | 8 | 0.002 | - |
| | Lean-control | FVT-SDT | 8 | 8 | 0.00931 | - |
| | Lean-control | FVT-PyT | 8 | 8 | 0.00931 | - |
| | Lean-control | Unmodified-FVT | 8 | 8 | 0.002 | - |
| **t60** | FVT-ChP | Obese-control | 8 | 8 | 0.442 | 0.673 |
| | FVT-SDT | Obese-control | 8 | 8 | 0.674 | 0.674 |
| | FVT-PyT | Obese-control | 8 | 8 | 0.027 | 0.108 |
| | Unmodified-FVT | Obese-control | 8 | 8 | 0.505 | 0.673 |
| | Obese-control | Lean-control | 8 | 8 | 0.01 | - |
| | Lean-control | FVT-ChP | 8 | 8 | 0.045 | - |
| | Lean-control | FVT-SDT | 8 | 8 | 0.000931 | - |
| | Lean-control | FVT-PyT | 8 | 8 | 0.003 | - |
| | Lean-control | Unmodified-FVT | 8 | 8 | 0.002 | - |
| **t90** | FVT-ChP | Obese-control | 8 | 8 | 0.958 | 1 |
| | FVT-SDT | Obese-control | 8 | 8 | 1 | 1 |

| | | | | | | |
|---|---|---|---|---|---|---|
| | FVT-PyT | Obese-control | 8 | 8 | 0.066 | 0.264 |
| | Unmodified-FVT | Obese-control | 8 | 8 | 0.528 | 1 |
| | Obese-control | Lean-control | 8 | 8 | 0.013 | - |
| | Lean-control | FVT-ChP | 8 | 8 | 0.035 | - |
| | Lean-control | FVT-SDT | 8 | 8 | 0.000155 | - |
| | Lean-control | FVT-PyT | 8 | 8 | 0.004 | - |
| | Lean-control | Unmodified-FVT | 8 | 8 | 0.002 | - |
| | FVT-ChP | Obese-control | 8 | 8 | 0.04 | 0.16 |
| | FVT-SDT | Obese-control | 8 | 8 | 0.206 | 0.275 |
| | FVT-PyT | Obese-control | 8 | 8 | 0.141 | 0.275 |
| | Unmodified-FVT | Obese-control | 8 | 8 | 0.528 | 0.528 |
| **t120** | Obese-control | Lean-control | 8 | 8 | 0.006 | - |
| | Lean-control | FVT-ChP | 8 | 8 | 0.007 | - |
| | Lean-control | FVT-SDT | 8 | 8 | 0.002 | - |
| | Lean-control | FVT-PyT | 8 | 8 | 0.016 | - |
| | Lean-control | Unmodified-FVT | 8 | 8 | 0.003 | - |

Table S2: Overview of the oral glucose tolerance (OGTT) statistics at each time point at study week 18 (23 weeks old). Comparison was done with Wilcoxon rank-sum test (two-side) with FDR correction which was only conducted between the FVT treatments and obese control mice.

| | Group1 | Group2 | n1 | n2 | p | p.adj |
|---|---|---|---|---|---|---|
| **t0** | FVT-ChP | Obese-control | 8 | 8 | 0.712 | 0.713 |
| | FVT-SDT | Obese-control | 8 | 8 | 0.127 | 0.508 |
| | FVT-PyT | Obese-control | 8 | 8 | 0.713 | 0.713 |
| | Unmodified-FVT | Obese-control | 8 | 8 | 0.293 | 0.586 |
| | Obese-control | Lean-control | 8 | 8 | 0.004 | - |
| | Lean-control | FVT-ChP | 8 | 8 | 0.001 | - |
| | Lean-control | FVT-SDT | 8 | 8 | 0.013 | - |
| | Lean-control | FVT-PyT | 8 | 8 | 0.001 | - |
| | Lean-control | Unmodified-FVT | 8 | 8 | 0.092 | - |
| **t15** | FVT-ChP | Obese-control | 8 | 8 | 0.752 | 1 |
| | FVT-SDT | Obese-control | 8 | 8 | 0.141 | 0.564 |
| | FVT-PyT | Obese-control | 8 | 8 | 1 | 1 |
| | Unmodified-FVT | Obese-control | 8 | 8 | 1 | 1 |
| | Obese-control | Lean-control | 8 | 8 | 0.005 | - |
| | Lean-control | FVT-ChP | 8 | 8 | 0.000155 | - |
| | Lean-control | FVT-SDT | 8 | 8 | 0.000311 | - |
| | Lean-control | FVT-PyT | 8 | 8 | 0.001 | - |
| | Lean-control | Unmodified-FVT | 8 | 8 | 0.021 | - |
| **t30** | FVT-ChP | Obese-control | 8 | 8 | 0.878 | 0.878 |
| | FVT-SDT | Obese-control | 8 | 8 | 0.328 | 0.878 |
| | FVT-PyT | Obese-control | 8 | 8 | 0.721 | 0.878 |
| | Unmodified-FVT | Obese-control | 8 | 8 | 0.636 | 0.878 |
| | Obese-control | Lean-control | 8 | 8 | 0.002 | - |
| | Lean-control | FVT-ChP | 8 | 8 | 0.00923 | - |
| | Lean-control | FVT-SDT | 8 | 8 | 0.003 | - |
| | Lean-control | FVT-PyT | 8 | 8 | 0.003 | - |
| | Lean-control | Unmodified-FVT | 8 | 8 | 0.103 | - |
| **t60** | FVT-ChP | Obese-control | 8 | 8 | 0.021 | 0.083 |
| | FVT-SDT | Obese-control | 8 | 8 | 0.462 | 0.799 |
| | FVT-PyT | Obese-control | 8 | 8 | 0.875 | 0.875 |
| | Unmodified-FVT | Obese-control | 8 | 8 | 0.599 | 0.799 |
| | Obese-control | Lean-control | 8 | 8 | 0.000931 | - |
| | Lean-control | FVT-ChP | 8 | 8 | 0.007 | - |
| | Lean-control | FVT-SDT | 8 | 8 | 0.007 | - |
| | Lean-control | FVT-PyT | 8 | 8 | 0.002 | - |
| | Lean-control | Unmodified-FVT | 8 | 8 | 0.429 | - |
| **t90** | FVT-ChP | Obese-control | 8 | 8 | 0.014 | 0.054 |
| | FVT-SDT | Obese-control | 8 | 8 | 0.721 | 0.961 |

| | | | | | | |
|---|---|---|---|---|---|---|
| | FVT-PyT | Obese-control | 8 | 8 | 0.43 | 0.86 |
| | Unmodified-FVT | Obese-control | 8 | 8 | 1 | 1 |
| | Obese-control | Lean-control | 8 | 8 | 0.000155 | - |
| | Lean-control | FVT-ChP | 8 | 8 | 0.000915 | - |
| | Lean-control | FVT-SDT | 8 | 8 | 0.000923 | - |
| | Lean-control | FVT-PyT | 8 | 8 | 0.001 | - |
| | Lean-control | Unmodified-FVT | 8 | 8 | 1 | - |
| | FVT-ChP | Obese-control | 8 | 8 | 0.171 | 0.342 |
| | FVT-SDT | Obese-control | 8 | 8 | 0.269 | 0.359 |
| | FVT-PyT | Obese-control | 8 | 8 | 0.114 | 0.342 |
| | Unmodified-FVT | Obese-control | 8 | 8 | 0.874 | 0.874 |
| **t120** | Obese-control | Lean-control | 8 | 8 | 0.002 | - |
| | Lean-control | FVT-ChP | 8 | 8 | 0.000915 | - |
| | Lean-control | FVT-SDT | 8 | 8 | 0.000899 | - |
| | Lean-control | FVT-PyT | 8 | 8 | 0.005 | - |
| | Lean-control | Unmodified-FVT | 8 | 8 | 0.528 | - |

94 Table S3: List included bacterial and phage strains, relevant strain-specific information, and growth conditions.

| **Phage** | **DSM ID** | **Family** | **Genome type** | **Bacterial host** | **Strain ID** | **DSM ID** | **Incubation temperature (°C)** | **Media** |
|---|---|---|---|---|---|---|---|---|
| C2 | In house | *Ceduovirus c2* | dsDNA | *Lactococcus lactis* | MG1363 | DSM 4366 | 30 | M17 |
| T4 | 4505 | *Straboviridae* | dsDNA | *Escherichia coli* | Luria | DSM 613 | 37 | LB |
| PhiX174 | 4497 | *Microviridae* | ssDNA | *Escherichia coli* | PC 0886 | DSM 13127 | 37 | BHI |
| MS2 | 13767 | *Fiersviridae* | ssRNA | *Escherichia coli* | W1485 | DSM 5695 | 37 | NZCYM + 2mg/L streptomycin |
| Phi6 | 21518 | *Cystoviridae* | dsRNA | *Pseudomonas sp.* | HER 1102 | DSM 21482 | 25 | TSB |

95

Table S4: Overview of cage-associated effects of phenotypical factors statistics. Comparison was done with Wilcoxon rank-sum test (two-side).

**FVT-ChP**

| Phenotypical factors | Cage | Cage | n1 | n2 | statistic | p |
|---|---|---|---|---|---|---|
| ewat_mg | 6 | 12 | 4 | 4 | 5 | 0.486 |
| Weight_gain_perc | 6 | 12 | 4 | 4 | 12 | 0.343 |
| Cumulative_score | 6 | 12 | 4 | 4 | 8 | 1 |
| Steatosis | 6 | 12 | 4 | 4 | 8 | 1 |
| Inflammation | - | - | - | - | - | - |
| Hepatocyte_injury | 6 | 12 | 4 | 4 | 8 | 1 |
| OGTT_t0_w18 | 6 | 12 | 4 | 4 | 11 | 0.486 |
| OGTT_t15_w18 | 6 | 12 | 4 | 4 | 12 | 0.343 |
| OGTT_t30_w18 | 6 | 12 | 4 | 4 | 12 | 0.343 |
| OGTT_t60_w18 | 6 | 12 | 4 | 4 | 10 | 0.686 |
| OGTT_t90_w18 | 6 | 12 | 4 | 4 | 9.5 | 0.772 |
| OGTT_t120_w18 | 6 | 12 | 4 | 4 | 14 | 0.114 |
| AUC1_w18 | 6 | 12 | 3 | 4 | 8 | 0.593 |
| OGTT_t0_w13 | 6 | 12 | 4 | 4 | 10 | 0.663 |
| OGTT_t15_w13 | 6 | 12 | 4 | 4 | 10 | 0.686 |
| OGTT_t30_w13 | 6 | 12 | 4 | 4 | 10.5 | 0.561 |
| OGTT_t60_w13 | 6 | 12 | 4 | 4 | 9 | 0.886 |
| OGTT_t90_w13 | 6 | 12 | 4 | 4 | 6 | 0.663 |
| OGTT_t120_w13 | 6 | 12 | 4 | 4 | 10.5 | 0.559 |
| AUC1_w13 | 6 | 12 | 4 | 3 | 9 | 0.4 |
| fat_Dendritic_cells | 6 | 12 | 4 | 3 | 9 | 0.4 |
| fat_Macrophages | 6 | 12 | 4 | 3 | 11 | 0.114 |
| fat_M1_macrophages | 6 | 12 | 4 | 3 | 1 | 0.114 |
| fat_B_cells | 6 | 12 | 4 | 3 | 5 | 0.857 |
| fat_Cytotoxic_T_cells | 6 | 12 | 4 | 3 | 8 | 0.593 |
| fat_Th_cells | 6 | 12 | 4 | 3 | 9 | 0.4 |
| fat_Central_memory_T_cells_CD8 | 6 | 12 | 4 | 3 | 3 | 0.4 |
| fat_Effector_memory_T_cells_CD8 | 6 | 12 | 4 | 3 | 4 | 0.629 |
| fat_Naive_T_cells_CD8 | 6 | 12 | 4 | 3 | 8 | 0.629 |
| fat_Central_memory_T_cells_CD4 | 6 | 12 | 4 | 3 | 3 | 0.4 |
| fat_Effector_memory_T_cells_CD4 | 6 | 12 | 4 | 3 | 7 | 0.857 |
| fat_Naive_T_cells_CD4 | 6 | 12 | 4 | 3 | 4 | 0.629 |
| Bac: Shannon DI - Arrival | 12 | 6 | 3 | 3 | 0 | 0.1 |
| Bac: Shannon DI – Before $1^{st}$ FVT | 12 | 6 | 4 | 4 | 8 | 1 |
| Bac: Shannon DI – 1w after FVT | 12 | 6 | 4 | 3 | 4 | 0.629 |
| Bac: Shannon DI - Termination | 12 | 6 | 4 | 4 | 5 | 0.486 |
| Bac: Bray-Curtis - Arrival | 12 | 6 | 3 | 3 | 1.26 | 0.100 |
| Bac: Bray-Curtis – Before $1^{st}$ FVT | 12 | 6 | 4 | 4 | 1.53 | 0.089 |
| Bac: Bray-Curtis – 1w after FVT | 12 | 6 | 4 | 3 | 1.55 | 0.032 |
| Bac: Bray-Curtis - Termination | 12 | 6 | 4 | 4 | 1.66 | 0.037 |
| Vir: Shannon DI - Arrival | 12 | 6 | 4 | 4 | 7 | 0.886 |
| Vir: Shannon DI – 1w after FVT | 12 | 6 | 4 | 3 | 7 | 0.857 |
| Vir: Shannon DI - Termination | 12 | 6 | 4 | 4 | 11 | 0.486 |
| Vir: Bray-Curtis - Arrival | 12 | 6 | 4 | 4 | 0.87 | 0.52 |
| Vir: Bray-Curtis – 1w after FVT | 12 | 6 | 4 | 3 | 1.12 | 0.36 |
| Vir: Bray-Curtis - Termination | 12 | 6 | 4 | 4 | 1.57 | 0.139 |

**FVT-SDT**

| Phenotypical factors | Cage | Cage | n1 | n2 | statistic | p |
|---|---|---|---|---|---|---|
| ewat_mg | 5 | 11 | 4 | 4 | 0 | 0.0286 |
| Weight_gain_perc | 5 | 11 | 4 | 4 | 10 | 0.686 |
| Cumulative_score | 5 | 11 | 4 | 4 | 8 | 1 |
| Steatosis | - | - | - | - | - | - |
| Inflammation | - | - | - | - | - | - |
| Hepatocyte_injury | - | - | - | - | - | - |
| OGTT_t0_w18 | 5 | 11 | 4 | 4 | 6 | 0.663 |
| OGTT_t15_w18 | 5 | 11 | 4 | 4 | 4 | 0.343 |
| OGTT_t30_w18 | 5 | 11 | 4 | 4 | 5 | 0.486 |
| OGTT_t60_w18 | 5 | 11 | 4 | 4 | 2 | 0.114 |
| OGTT_t90_w18 | 5 | 11 | 4 | 4 | 0 | 0.0286 |
| OGTT_t120_w18 | 5 | 11 | 4 | 4 | 2 | 0.114 |
| AUC1_w18 | 5 | 11 | 4 | 3 | 2 | 0.229 |
| OGTT_t0_w13 | 5 | 11 | 4 | 4 | 16 | 0.0286 |
| OGTT_t15_w13 | 5 | 11 | 4 | 4 | 13 | 0.2 |
| OGTT_t30_w13 | 5 | 11 | 4 | 4 | 13 | 0.2 |
| OGTT_t60_w13 | 5 | 11 | 4 | 4 | 9.5 | 0.772 |
| OGTT_t90_w13 | 5 | 11 | 4 | 4 | 15 | 0.0571 |
| OGTT_t120_w13 | 5 | 11 | 4 | 4 | 13 | 0.183 |
| AUC1_w13 | 5 | 11 | 3 | 4 | 8 | 0.629 |
| fat_Dendritic_cells | 5 | 11 | 3 | 4 | 12 | 0.0571 |
| fat_Macrophages | 5 | 11 | 3 | 4 | 3 | 0.4 |
| fat_M1_macrophages | 5 | 11 | 3 | 4 | 8 | 0.629 |
| fat_B_cells | 5 | 11 | 3 | 4 | 9 | 0.4 |
| fat_Cytotoxic_T_cells | 5 | 11 | 3 | 4 | 9 | 0.4 |
| fat_Th_cells | 5 | 11 | 3 | 4 | 8 | 0.629 |
| fat_Central_memory_T_cells_CD8 | 5 | 11 | 3 | 4 | 5 | 0.857 |
| fat_Effector_memory_T_cells_CD8 | 5 | 11 | 3 | 4 | 9 | 0.4 |
| fat_Naive_T_cells_CD8 | 5 | 11 | 3 | 4 | 1 | 0.114 |
| fat_Central_memory_T_cells_CD4 | 5 | 11 | 3 | 4 | 8 | 0.629 |
| fat_Effector_memory_T_cells_CD4 | 5 | 11 | 3 | 4 | 6 | 1 |
| fat_Naive_T_cells_CD4 | 5 | 11 | 3 | 4 | 5 | 0.857 |
| Bac: Shannon DI - Arrival | 11 | 5 | 4 | 4 | 4 | 0.343 |
| Bac: Shannon DI – Before $1^{st}$ FVT | 11 | 5 | 4 | 4 | 6 | 0.686 |
| Bac: Shannon DI – 1w after FVT | 11 | 5 | 4 | 3 | 2 | 0.229 |
| Bac: Shannon DI - Termination | 11 | 5 | 4 | 3 | 6 | 1 |
| Bac: Bray-Curtis - Arrival | 11 | 5 | 4 | 4 | 1.46 | 0.161 |
| Bac: Bray-Curtis – Before $1^{st}$ FVT | 11 | 5 | 4 | 4 | 1.82 | 0.025 |
| Bac: Bray-Curtis – 1w after FVT | 11 | 5 | 4 | 3 | 1.40 | 0.124 |
| Bac: Bray-Curtis - Termination | 11 | 5 | 4 | 3 | 2.87 | 0.035 |
| Vir: Shannon DI - Arrival | 11 | 5 | 4 | 3 | 4 | 0.629 |
| Vir: Shannon DI – 1w after FVT | 11 | 5 | 3 | 4 | 6 | 1 |
| Vir: Shannon DI - Termination | 11 | 5 | 3 | 4 | 9 | 0.4 |
| Vir: Bray-Curtis - Arrival | 11 | 5 | 4 | 3 | 1.11 | 0.278 |
| Vir: Bray-Curtis – 1w after FVT | 11 | 5 | 3 | 4 | 1.78 | 0.027 |
| Vir: Bray-Curtis - Termination | 11 | 5 | 3 | 4 | 1.45 | 0.146 |

**FVT-PyT**

| Phenotypical factors | Cage | Cage | n1 | n2 | statistic | p |
|---|---|---|---|---|---|---|
| ewat_mg | 2 | 8 | 4 | 4 | 7 | 0.886 |
| Weight_gain_perc | 2 | 8 | 4 | 4 | 13 | 0.2 |
| Cumulative_score | 2 | 8 | 4 | 4 | 6.5 | 0.766 |
| Steatosis | 2 | 8 | 4 | 4 | 7 | 0.874 |
| Inflammation | 2 | 8 | 4 | 4 | 6 | 0.608 |
| Hepatocyte_injury | 2 | 8 | 4 | 4 | 7 | 0.874 |
| OGTT_t0_w18 | 2 | 8 | 4 | 4 | 9 | 0.886 |
| OGTT_t15_w18 | 2 | 8 | 4 | 4 | 3 | 0.2 |
| OGTT_t30_w18 | 2 | 8 | 4 | 4 | 4 | 0.343 |
| OGTT_t60_w18 | 2 | 8 | 4 | 4 | 2 | 0.114 |
| OGTT_t90_w18 | 2 | 8 | 4 | 4 | 2.5 | 0.146 |
| OGTT_t120_w18 | 2 | 8 | 4 | 4 | 4 | 0.309 |
| AUC1_w18 | 2 | 8 | 4 | 4 | 2 | 0.114 |
| OGTT_t0_w13 | 2 | 8 | 4 | 4 | 11 | 0.486 |
| OGTT_t15_w13 | 2 | 8 | 4 | 4 | 9 | 0.886 |
| OGTT_t30_w13 | 2 | 8 | 4 | 4 | 7 | 0.886 |
| OGTT_t60_w13 | 2 | 8 | 4 | 4 | 7 | 0.882 |
| OGTT_t90_w13 | 2 | 8 | 4 | 4 | 6 | 0.686 |
| OGTT_t120_w13 | 2 | 8 | 4 | 4 | 5 | 0.486 |
| AUC1_w13 | 2 | 8 | 4 | 4 | 7 | 0.886 |
| fat_Dendritic_cells | 2 | 8 | 4 | 4 | 8 | 1 |
| fat_Macrophages | 2 | 8 | 4 | 4 | 11 | 0.486 |
| fat_M1_macrophages | 2 | 8 | 4 | 4 | 6 | 0.686 |
| fat_B_cells | 2 | 8 | 4 | 4 | 7 | 0.886 |
| fat_Cytotoxic_T_cells | 2 | 8 | 4 | 4 | 14 | 0.114 |
| fat_Th_cells | 2 | 8 | 4 | 4 | 12 | 0.343 |

**Unmodified-FVT**

| Phenotypical factors | Cage | Cage | n1 | n2 | statistic | p |
|---|---|---|---|---|---|---|
| ewat_mg | 3 | 9 | 4 | 4 | 8 | 1 |
| Weight_gain_perc | 3 | 9 | 4 | 4 | 16 | 0.0286 |
| Cumulative_score | 3 | 9 | 4 | 4 | 8.5 | 1 |
| Steatosis | 3 | 9 | 4 | 4 | 10 | 0.642 |
| Inflammation | 3 | 9 | 4 | 4 | 6 | 0.608 |
| Hepatocyte_injury | 3 | 9 | 4 | 4 | 9 | 0.874 |
| OGTT_t0_w18 | 3 | 9 | 4 | 4 | 15 | 0.0571 |
| OGTT_t15_w18 | 3 | 9 | 4 | 4 | 12 | 0.343 |
| OGTT_t30_w18 | 3 | 9 | 4 | 4 | 13 | 0.2 |
| OGTT_t60_w18 | 3 | 9 | 4 | 4 | 11 | 0.486 |
| OGTT_t90_w18 | 3 | 9 | 4 | 4 | 16 | 0.0294 |
| OGTT_t120_w18 | 3 | 9 | 4 | 4 | 16 | 0.0286 |
| AUC1_w18 | 3 | 9 | 3 | 4 | 9 | 0.4 |
| OGTT_t0_w13 | 3 | 9 | 4 | 4 | 10 | 0.663 |
| OGTT_t15_w13 | 3 | 9 | 4 | 4 | 8 | 1 |
| OGTT_t30_w13 | 3 | 9 | 4 | 4 | 9 | 0.886 |
| OGTT_t60_w13 | 3 | 9 | 4 | 4 | 7 | 0.886 |
| OGTT_t90_w13 | 3 | 9 | 4 | 4 | 8 | 1 |
| OGTT_t120_w13 | 3 | 9 | 4 | 4 | 9 | 0.886 |
| AUC1_w13 | 3 | 9 | 4 | 4 | 8 | 1 |
| fat_Dendritic_cells | 3 | 9 | 4 | 4 | 10 | 0.686 |
| fat_Macrophages | 3 | 9 | 4 | 4 | 13 | 0.2 |
| fat_M1_macrophages | 3 | 9 | 4 | 4 | 9 | 0.886 |
| fat_B_cells | 3 | 9 | 4 | 4 | 9 | 0.886 |
| fat_Cytotoxic_T_cells | 3 | 9 | 4 | 4 | 12 | 0.343 |
| fat_Th_cells | 3 | 9 | 4 | 4 | 12 | 0.343 |

| | | | | | | | | | | | | | |
|---|---|---|---|---|---|---|---|---|---|---|---|---|---|
| fat_Central_memory_T_cells_CD8 | 2 | 8 | 4 | 4 | 7 | 0.886 | fat_Central_memory_T_cells_CD8 | 3 | 9 | 4 | 4 | 14 | 0.114 |
| fat_Effector_memory_T_cells_CD8 | 2 | 8 | 4 | 4 | 14 | 0.114 | fat_Effector_memory_T_cells_CD8 | 3 | 9 | 4 | 4 | 12 | 0.343 |
| fat_Naive_T_cells_CD8 | 2 | 8 | 4 | 4 | 2 | 0.114 | fat_Naive_T_cells_CD8 | 3 | 9 | 4 | 4 | 4 | 0.343 |
| fat_Central_memory_T_cells_CD4 | 2 | 8 | 4 | 4 | 5 | 0.486 | fat_Central_memory_T_cells_CD4 | 3 | 9 | 4 | 4 | 11 | 0.486 |
| fat_Effector_memory_T_cells_CD4 | 2 | 8 | 4 | 4 | 13 | 0.2 | fat_Effector_memory_T_cells_CD4 | 3 | 9 | 4 | 4 | 7 | 0.886 |
| fat_Naive_T_cells_CD4 | 2 | 8 | 4 | 4 | 2 | 0.114 | fat_Naive_T_cells_CD4 | 3 | 9 | 4 | 4 | 9 | 0.886 |
| Bac: Shannon DI - Arrival | 2 | 8 | 4 | 4 | 15 | 0.0571 | Bac: Shannon DI - Arrival | 3 | 9 | 4 | 3 | 4 | 0.629 |
| Bac: Shannon DI – Before 1st FVT | 2 | 8 | 4 | 4 | 13 | 0.2 | Bac: Shannon DI – Before 1st FVT | 3 | 9 | 4 | 4 | 7 | 0.886 |
| Bac: Shannon DI – 1w after FVT | 2 | 8 | 4 | 3 | 3 | 0.4 | Bac: Shannon DI – 1w after FVT | 3 | 9 | 3 | 2 | 1 | 0.4 |
| Bac: Shannon DI - Termination | 2 | 8 | 4 | 4 | 2 | 0.114 | Bac: Shannon DI - Termination | 3 | 9 | 4 | 4 | 4 | 0.343 |
| Bac: Bray-Curtis - Arrival | 2 | 8 | 4 | 4 | 1.82 | 0.027 | Bac: Bray-Curtis - Arrival | 3 | 9 | 4 | 3 | 3.13 | 0.036 |
| Bac: Bray-Curtis – Before 1st FVT | 2 | 8 | 4 | 4 | 2.09 | 0.02 | Bac: Bray-Curtis – Before 1st FVT | 3 | 9 | 4 | 4 | 1.93 | 0.024 |
| Bac: Bray-Curtis – 1w after FVT | 2 | 8 | 4 | 3 | 1.43 | 0.067 | Bac: Bray-Curtis – 1w after FVT | 3 | 9 | 3 | 2 | 1.37 | 0.1 |
| Bac: Bray-Curtis - Termination | 2 | 8 | 4 | 4 | 2.34 | 0.027 | Bac: Bray-Curtis - Termination | 3 | 9 | 4 | 4 | 2.00 | 0.049 |
| Vir: Shannon DI - Arrival | 2 | 8 | 4 | 4 | 10 | 0.686 | Vir: Shannon DI - Arrival | 3 | 9 | 4 | 4 | 11 | 0.486 |
| Vir: Shannon DI – 1w after FVT | 2 | 8 | 4 | 3 | 4 | 0.629 | Vir: Shannon DI – 1w after FVT | 3 | 9 | 3 | 2 | 3 | 1 |
| Vir: Shannon DI - Termination | 2 | 8 | 4 | 4 | 9 | 0.886 | Vir: Shannon DI - Termination | 3 | 9 | 4 | 4 | 9 | 0.886 |
| Vir: Bray-Curtis - Arrival | 2 | 8 | 4 | 4 | 2.79 | 0.022 | Vir: Bray-Curtis - Arrival | 3 | 9 | 4 | 4 | 2.62 | 0.028 |
| Vir: Bray-Curtis – 1w after FVT | 2 | 8 | 4 | 3 | 1.52 | 0.147 | Vir: Bray-Curtis – 1w after FVT | 3 | 9 | 3 | 2 | 3.13 | 0.1 |
| Vir: Bray-Curtis - Termination | 2 | 8 | 4 | 4 | 1.28 | 0.246 | Vir: Bray-Curtis - Termination | 3 | 9 | 4 | 4 | 1.55 | 0.179 |

| **Obese-control** | | | | | | | **Lean-control** | | | | | | |
|---|---|---|---|---|---|---|---|---|---|---|---|---|---|
| **Phenotypical factors** | **Cage** | **Cage** | **n1** | **n2** | **statistic** | **p** | **Phenotypical factors** | **Cage** | **Cage** | **n1** | **n2** | **statistic** | **p** |
| ewat_mg | 4 | 10 | 4 | 4 | 3 | 0.2 | ewat_mg | 1 | 7 | 4 | 4 | 2 | 0.114 |
| Weight_gain_perc | 4 | 10 | 4 | 4 | 16 | 0.0286 | Weight_gain_perc | 1 | 7 | 4 | 4 | 10 | 0.686 |
| Cumulative_score | 4 | 10 | 4 | 4 | 9 | 0.874 | Cumulative_score | 1 | 7 | 4 | 4 | 6 | 0.655 |
| Steatosis | 4 | 10 | 4 | 4 | 12 | 0.181 | Steatosis | 1 | 7 | 4 | 4 | 3 | 0.161 |
| Inflammation | 4 | 10 | 4 | 4 | 4 | 0.181 | Inflammation | 1 | 7 | 4 | 4 | 8 | 1 |
| Hepatocyte_injury | 4 | 10 | 4 | 4 | 10 | 0.453 | Hepatocyte_injury | 1 | 7 | 4 | 4 | 10 | 0.608 |
| OGTT_t0_w18 | 4 | 10 | 4 | 4 | 5 | 0.465 | OGTT_t0_w18 | 1 | 7 | 4 | 4 | 8 | 1 |
| OGTT_t15_w18 | 4 | 10 | 4 | 4 | 6.5 | 0.772 | OGTT_t15_w18 | 1 | 7 | 4 | 4 | 12 | 0.343 |
| OGTT_t30_w18 | 4 | 10 | 4 | 4 | 10 | 0.686 | OGTT_t30_w18 | 1 | 7 | 4 | 4 | 14.5 | 0.0814 |
| OGTT_t60_w18 | 4 | 10 | 4 | 4 | 13 | 0.2 | OGTT_t60_w18 | 1 | 7 | 4 | 4 | 6 | 0.663 |
| OGTT_t90_w18 | 4 | 10 | 4 | 4 | 13 | 0.2 | OGTT_t90_w18 | 1 | 7 | 4 | 4 | 4 | 0.343 |
| OGTT_t120_w18 | 4 | 10 | 4 | 4 | 14 | 0.104 | OGTT_t120_w18 | 1 | 7 | 4 | 4 | 10 | 0.663 |
| AUC1_w18 | 4 | 10 | 4 | 4 | 12 | 0.343 | AUC1_w18 | 1 | 7 | 4 | 4 | 9 | 0.886 |
| OGTT_t0_w13 | 4 | 10 | 4 | 4 | 13.5 | 0.146 | OGTT_t0_w13 | 1 | 7 | 4 | 4 | 6.5 | 0.772 |
| OGTT_t15_w13 | 4 | 10 | 4 | 4 | 5 | 0.486 | OGTT_t15_w13 | 1 | 7 | 4 | 4 | 10 | 0.686 |
| OGTT_t30_w13 | 4 | 10 | 4 | 4 | 6 | 0.686 | OGTT_t30_w13 | 1 | 7 | 4 | 4 | 6.5 | 0.772 |
| OGTT_t60_w13 | 4 | 10 | 4 | 4 | 5 | 0.486 | OGTT_t60_w13 | 1 | 7 | 4 | 4 | 13.5 | 0.146 |
| OGTT_t90_w13 | 4 | 10 | 4 | 4 | 8.5 | 1 | OGTT_t90_w13 | 1 | 7 | 4 | 4 | 11 | 0.465 |
| OGTT_t120_w13 | 4 | 10 | 4 | 4 | 9 | 0.885 | OGTT_t120_w13 | 1 | 7 | 4 | 4 | 0 | 0.0294 |
| AUC1_w13 | 4 | 10 | 4 | 4 | 6 | 0.686 | AUC1_w13 | 1 | 7 | 4 | 4 | 13 | 0.2 |
| fat_Dendritic_cells | 4 | 10 | 4 | 4 | 6 | 0.686 | fat_Dendritic_cells | 1 | 7 | 2 | 4 | 0 | 0.133 |
| fat_Macrophages | 4 | 10 | 4 | 4 | 16 | 0.0286 | fat_Macrophages | 1 | 7 | 2 | 4 | 8 | 0.133 |
| fat_M1_macrophages | 4 | 10 | 4 | 4 | 2 | 0.114 | fat_M1_macrophages | 1 | 7 | 2 | 4 | 4 | 1 |
| fat_B_cells | 4 | 10 | 4 | 4 | 4 | 0.343 | fat_B_cells | 1 | 7 | 2 | 4 | 5 | 0.8 |
| fat_Cytotoxic_T_cells | 4 | 10 | 4 | 4 | 8 | 1 | fat_Cytotoxic_T_cells | 1 | 7 | 3 | 4 | 4 | 0.629 |
| fat_Th_cells | 4 | 10 | 4 | 4 | 16 | 0.0286 | fat_Th_cells | 1 | 7 | 3 | 4 | 8 | 0.629 |
| fat_Central_memory_T_cells_CD8 | 4 | 10 | 4 | 4 | 16 | 0.0286 | fat_Central_memory_T_cells_CD8 | 1 | 7 | 3 | 4 | 9 | 0.4 |
| fat_Effector_memory_T_cells_CD8 | 4 | 10 | 4 | 4 | 8 | 1 | fat_Effector_memory_T_cells_CD8 | 1 | 7 | 3 | 4 | 0 | 0.0571 |
| fat_Naive_T_cells_CD8 | 4 | 10 | 4 | 4 | 1 | 0.0571 | fat_Naive_T_cells_CD8 | 1 | 7 | 3 | 4 | 12 | 0.0571 |
| fat_Central_memory_T_cells_CD4 | 4 | 10 | 4 | 4 | 5 | 0.486 | fat_Central_memory_T_cells_CD4 | 1 | 7 | 3 | 4 | 4 | 0.629 |
| fat_Effector_memory_T_cells_CD4 | 4 | 10 | 4 | 4 | 11 | 0.486 | fat_Effector_memory_T_cells_CD4 | 1 | 7 | 3 | 4 | 9 | 0.4 |
| fat_Naive_T_cells_CD4 | 4 | 10 | 4 | 4 | 5 | 0.486 | fat_Naive_T_cells_CD4 | 1 | 7 | 3 | 4 | 1 | 0.114 |
| Bac: Shannon DI - Arrival | 10 | 4 | 4 | 4 | 7 | 0.886 | Bac: Shannon DI - Arrival | 1 | 7 | 4 | 4 | 7 | 0.886 |
| Bac: Shannon DI – Before 1st FVT | 10 | 4 | 4 | 4 | 12 | 0.343 | Bac: Shannon DI – Before 1st FVT | 1 | 7 | 4 | 4 | 11 | 0.486 |
| Bac: Shannon DI – 1w after FVT | 10 | 4 | 3 | 4 | 5 | 0.857 | Bac: Shannon DI – 1w after FVT | 1 | 7 | 4 | 4 | 7 | 0.886 |
| Bac: Shannon DI - Termination | 10 | 4 | 4 | 4 | 6 | 0.686 | Bac: Shannon DI - Termination | 1 | 7 | 4 | 4 | 1 | 0.0571 |
| sBac: Bray-Curtis - Arrival | 10 | 4 | 4 | 4 | 2.00 | 0.028 | Bac: Bray-Curtis - Arrival | 1 | 7 | 4 | 4 | 2.35 | 0.026 |
| Bac: Bray-Curtis – Before 1st FVT | 10 | 4 | 4 | 4 | 2.35 | 0.026 | Bac: Bray-Curtis – Before 1st FVT | 1 | 7 | 4 | 4 | 2.61 | 0.031 |
| Bac: Bray-Curtis – 1w after FVT | 10 | 4 | 3 | 4 | 1.59 | 0.035 | Bac: Bray-Curtis – 1w after FVT | 1 | 7 | 4 | 4 | 1.98 | 0.022 |
| Bac: Bray-Curtis - Termination | 10 | 4 | 4 | 4 | 2.27 | 0.034 | Bac: Bray-Curtis - Termination | 1 | 7 | 4 | 4 | 3.57 | 0.024 |
| Vir: Shannon DI - Arrival | 10 | 4 | 4 | 4 | 8 | 1 | Vir: Shannon DI - Arrival | 1 | 7 | 4 | 3 | 2 | 0.229 |
| Vir: Shannon DI – 1w after FVT | 10 | 4 | 3 | 4 | 6 | 1 | Vir: Shannon DI – 1w after FVT | 1 | 7 | 3 | 4 | 6 | 1 |
| Vir: Shannon DI - Termination | 10 | 4 | 4 | 4 | 8 | 1 | Vir: Shannon DI - Termination | 1 | 7 | 3 | 4 | 5 | 0.857 |
| Vir: Bray-Curtis - Arrival | 10 | 4 | 4 | 4 | 1.78 | 0.033 | Vir: Bray-Curtis - Arrival | 1 | 7 | 4 | 3 | 3.21 | 0.004 |
| Vir: Bray-Curtis – 1w after FVT | 10 | 4 | 3 | 4 | 2.07 | 0.025 | Vir: Bray-Curtis – 1w after FVT | 1 | 7 | 3 | 4 | 0.91 | 0.531 |
| Vir: Bray-Curtis - Termination | 10 | 4 | 4 | 4 | 2.47 | 0.033 | Vir: Bray-Curtis - Termination | 1 | 7 | 3 | 4 | 2.45 | 0.064 |
